# Chromosome-scale reference genome and RAD-based genetic map of yellow starthistle (*Centaurea solstitialis*) reveal putative structural variation and QTL associated with invader traits

**DOI:** 10.1101/2022.09.28.509992

**Authors:** Bryan Reatini, Jessie A. Pelosi, F. Alice Cang, Qiuyu Jiang, Michael T. W. McKibben, Michael S. Barker, Loren H. Rieseberg, Katrina M. Dlugosch

**Author notes:** Author for Correspondence: Bryan Reatini, Department of Ecology and Evolutionary Biology, University of Arizona, Tucson, Arizona.

## Abstract

Invasive species offer outstanding opportunities to identify the genomic sources of variation that contribute to rapid adaptation, as well as the genetic mechanisms facilitating invasions. The Eurasian plant yellow starthistle (*Centaurea solstitialis*) is highly invasive in North and South American grasslands and known to have evolved increased growth and reproduction during invasion. Here we develop new genomic resources for *C. solstitialis* and map the genetic basis of invasiveness traits. We present a chromosome-scale (1N = 8) reference genome using PacBio CLR and Dovetail Omni-C technologies, and functional gene annotation using RNAseq. We find repeat structure typical of the family Asteraceae, with over 25% of gene content derived from ancestral whole genome duplications (paleologs). Using an F2 mapping population derived from a cross between native and invading parents, with a restriction site-associated DNA (RAD)-based genetic map, we validate the assembly and identify 13 QTL underpinning size traits that have evolved during invasion. We find evidence that large effect QTL may be associated with structural variants between native and invading genotypes, including a variant with an overdominant and pleiotropic effect on key invader traits. We also find evidence of significant paleolog enrichment under two QTL. Our results add to growing evidence of the importance of structural variants in evolution, and to understanding of the rapid evolution of invaders.

**Significance Statement:** Invasive species often evolve rapidly in new environments, potentially informing our understanding of the genomic basis of adaptation, but genomic studies of these non-model systems are limited. We provide a chromosome-scale reference genome, annotation, and genetic map for the invasive plant yellow starthistle, and we investigate the genetic basis of invader trait evolution in this system. We find regions of the genome with large effects on traits that differ between native and invading genotypes, and evidence suggesting genome structural variants and past genome duplications could play a role in rapid adaptation of invading populations. These genomic resources and evolutionary insights aid in our understanding of the sources of genomic variation for adaptation, and how their evolution facilitates invasion.

## Introduction

Biological invasions provide unique opportunities to study how rapid genome evolution can contribute to population establishment, growth, and range expansion in novel environments (McGaughran et al. 2024). Indeed, what types of genomic variation will fuel rapid adaptation has been a longstanding question in evolution (Orr 1998), and invading populations are increasingly providing insights into the adaptive role of different classes of genomic variants (reviewed in Dlugosch, Anderson, et al. 2015; McGaughran et al. 2024; C. E. Lee 2002). For instance, chromosomal inversions have been identified underpinning climate adaptation in invading *Ambrosia artemisiifolia* (Battlay et al. 2022) and transposable element insertions have been identified underpinning adaptive shifts in flowering time in invading *Capsella rubella* (Niu et al. 2019), exemplifying how structural variation can play an important role in the evolution of high fitness genotypes. Whole genome duplications - either in the form of autopolyploidy or allopolyploidy - have also been found to promote invasion success by, for example, generating genetic variation to fuel adaptive evolution, increasing functional plasticity, and providing redundant gene duplicates that can lead to evolutionary novelty via neofunctionalization, even many millions of years after the polyploidy event (te Beest et al. 2012; Mounger et al. 2021; Qi et al. 2021).

Genomic analyses of invasions are also providing opportunities to learn how and when invasions occur, and to identify opportunities for management. There is longstanding interest in the role of genetic variation and evolution in facilitating the establishment and invasion of introduced species (Dlugosch, Anderson, et al. 2015). Research in this area is revealing when and how population bottlenecks might affect fitness of founding populations (Nei, Maruyama, and Chakraborty 1975; Estoup et al. 2016; Peischl et al. 2018), the importance of genomic admixture (B. S. Barker et al. 2019; Reatini and Vision 2020), and the genomic pathways that might underlie invader adaptations and potential sources of control, such as those involved in enemy interactions (Battlay et al. 2022). Given the variety of ways genome evolution could contribute to rapid adaptation, comprehensive investigations of invasion genomics require high-quality genomic resources, including annotations of genes, gene duplicates, and repetitive elements, as well as assembly of complete reference genomes that allow for the identification of structural variants. The development of such resources for an increasingly broad array of invaders will inform our understanding of both how genomes evolve in wild populations, and how invasions occur and can be managed (Dlugosch, Anderson, et al. 2015; Bock et al. 2015; McGaughran et al. 2024).

Here we establish a reference genome and annotations for the highly invasive plant yellow starthistle, *Centaurea solstitialis* L. (Asteraceae), and we map genomic regions associated with invader traits known to have evolved in this species. *Centaurea solstitialis* is invasive on at least four continents and occupies a broad distribution throughout its native range in Eurasia (Maddox, Mayfield, and Poritz 1985). The species was introduced from western Europe to South America in the 1600s and then to North America in the 1800s, and has since spread aggressively as a noxious weed of grasslands in the western United States and Argentina (B. S. Barker et al. 2017; Gerlach 1997). In the United States, plant traits in the severe invasion of California have been well-studied, and invaders are known to have evolved an increase in plant size (B. S. Barker et al. 2017; Widmer et al. 2007; Dlugosch, Cang, et al. 2015; Eriksen et al. 2012; Montesinos and Callaway 2018). Larger plant size is associated with increase reproduction (Dlugosch, Cang, et al. 2015) and competitive ability (Montesinos and Callaway 2017; Montesinos, Graebner, and Callaway 2019), and is predicted to lead to higher population growth rates for invader genotypes relative to native genotypes (Dlugosch, Cang, et al. 2015).

How *C. solstitialis* has rapidly evolved invasiveness, and the genomic mechanisms involved in achieving larger size, are not yet known, but the system is well suited to genomic analyses. The species is annual, diploid (1N = 8; Widmer et al. 2007), and obligately outcrossing, with a modest genome size of 840 Mbp (Irimia et al. 2017; Bancheva and Greilhuber 2006; Heiser Jr. and Whitaker 1948; Cang 2024). Population genomic studies have determined that the invading California lineage has evolved from a single native range source in western Europe, which provides an ancestral comparison for the evolution of the invaders (B. S. Barker et al. 2017). Finally, the evolutionary history of *C. solstitialis* includes an ancestral whole genome duplication at the base of the Asteraceae (M. S. Barker et al. 2008; 2016), providing an opportunity to identify the contribution of this event to contemporary genomic variation and rapid evolution (Qi et al. 2021).

We present a chromosome-scale reference genome for an outbred wild individual of *C. solstitialis* collected from Canales, Spain - the western European native source population of the Californian invasion. We characterize the content and structure of the *C. solstitialis* genome using annotations of repetitive elements, functional genes, and gene duplicates including paleologs (gene duplicates derived from ancestral whole genome duplications), and gene synteny comparisons between starthistle and other representatives of *Asteraceae*. We validate the reference genome assembly by comparison with a genetic map constructed from an F2 mapping population. We then leverage these genomic resources to identify quantitative trait loci (QTL) underpinning increased plant size in the Californian populations, identifying candidate genes within those QTL regions, and uncovering evidence suggesting that structural variants and paleologs may be forms of genomic variation contributing to evolution at plant size QTL. Finally, we also find that large scale genome rearrangements have characterized chromosome evolution across longer timescales within the Carduoideae subfamily of Asteraceae. Together, our findings contribute to growing evidence of the importance of structural variation in standing variation, and lay a foundation for investigating the contribution of genome evolution to the invasiveness of *C. solstitialis*, and to genome evolution more broadly within the thistle subfamily of Asteraceae.

## Results

### Genome assembly and annotation

We constructed a chromosome-scale reference genome assembly for *C. solstitialis* using a combination of PacBio CLR and Dovetail Omni-C sequencing approaches. The final assembly was 746Mb in length and contained 2,969 contigs (1.3Mb contig N50) which were anchored on 1,080 scaffolds; eight primary scaffolds comprised 94.8% of the total sequence content of the assembly (L90=8, Figure 1A), and matched the haploid chromosome number of *C. solstitialis* (1N = 8; Widmer et al. 2007). These eight scaffolds totalled 725.4 Mbp, which is 86.3% of the 840 Mbp average genome size reported for *C. solstitialis*, as estimated by flow cytometry (Cang 2024). An additional 1,072 unplaced scaffolds totaled 39.6 Mbp. Of the 2,326 conserved single copy orthologs (BUSCOs) in the eudicot_odb10 database, 2,106 (93.38% were found in the eight primary scaffolds: 1,917 (86.63%) were complete and single-copy and 189 (6.75%) were complete and duplicated (Table 1). Only two of the BUSCOs that were missing in the eight primary scaffolds were located in the trailing scaffolds. In addition, 20 of the BUSCOs that were found in single-copy in the eight primary scaffolds were duplicated in the trailing scaffolds, suggesting that some trailing scaffolds possibly included haplotype variants of the primary scaffolds. Given their nearly complete coverage of the genome, we hereafter refer to the eight primary scaffolds as chromosomes 1-8, numbered by descending size.

**Figure 1.**
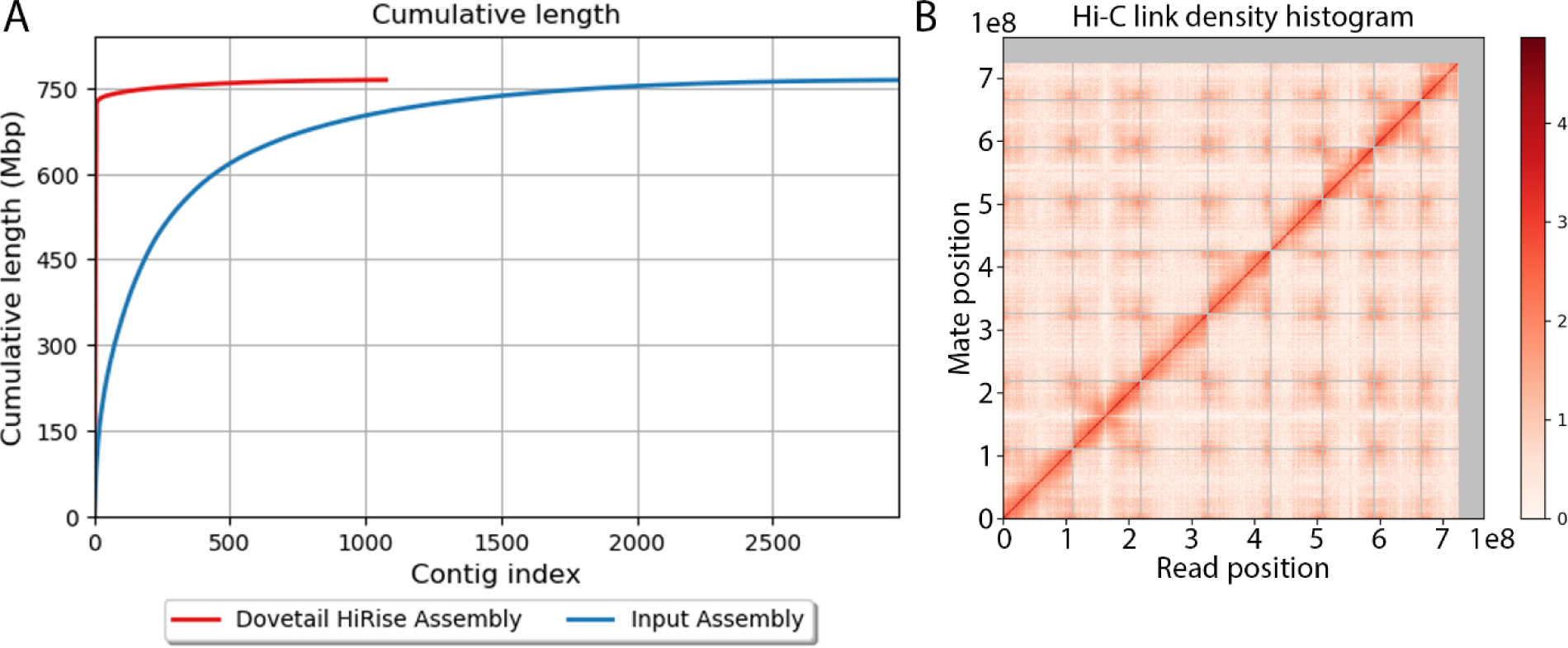
Scaffolding to the chromosome-level of the *C. solstitialis* genome using Omni-C data via the dovetail HiRise pipeline. A) Cumulative length of scaffolds in the final HiRise assembly versus the initial PacBio CLR assembly. B) Link density of read pairs from the Dovetail Omni-C data mapped to the final *C. solstitialis* assembly. X and Y-axes represent the mapping positions of the first and second read in each read pair, respectively. Positions are binned, and link density within bins are represented as a heat map. Grey bars denote boundaries between scaffolds.

**Table 1.** *Centaurea solstitialis* genome assembly and annotation statistics.

| Metric | Final Omni-C Assembly |
| --- | --- |
| **Assembly** |  |
| **Total length** | 765,086,668 |
| **Total number of contigs** | 2,969 |
| **Total number of scaffolds** | 1,080 |
| **Contig N50** | 1,356,249 |
| **Scaffold N50** | 100,696,929 |
| **Scaffold L90** | 8 |
| **Complete BUSCOs** | 2172 (94.37%) |
| **Complete and single copy BUSCOs** | 2015 (86.63%) |
| **Annotation** |  |
| **Total repetitive DNA** | 63.31% |
| **Transposable elements** | 31.1% |
| **Simple repeats** | 1.16% |
| **Total number of genes** | 34,323 |
| **Total coding region (bp)** | 59,845,476 |
| **Average gene length (bp)** | 1,743 |

We characterized the content of the genome in terms of its repetitive elements, functional regions, paleologs, centromere locations, and chromosome structure. An estimated 63.3% of the genome was repetitive DNA, with 29.04% identified as class I transposable elements (retrotransposons) and 2.07% identified as class II transposable elements (DNA transposons) (Table 1). A total of 481,954 retroelements - composing 12% of the genome - were successfully classified using the high-confidence nrTEplants database, and included 122,163 Ty1/Copia and 172,604 Gypsy long terminal repeat retrotransposons (LTR-RTs).

We identified functional regions of the genome using RNAseq of three native and three invading individuals, including the individual sequenced for the reference genome. The complete reference annotation included 34,323 predicted gene models encompassing a total of 59.8 Mbp of the reference sequence (Table 1). Of the 32,431 genes that fell on the eight putative chromosomes, 74% (24,011) were able to be functionally annotated using the UniProt database. A total of 616 tRNA prediction models were identified using tRNAscan-SE. Using the cumulative frequency of annotation edit distance (with 0 being perfect support and 1 being no support) as a metric of annotation quality, the majority of genes in the complete annotation were well supported by overlapping aligned RNAseq and protein homology data, with 80% of all genes having AED scores of <0.5 (Supplemental Figure 1).

Paleologs from ancestral genome duplications may persist in blocks that are identifiable despite fractionation over time (Cheng et al. 2018). The *C. solstitialis* lineage most recently experienced a putative hexaploidy in its ancestry at the base of the family, Asteraceae (M. S. Barker et al. 2016; 2008), and we identified blocks of paleologs using syntenic comparisons with an outgroup (carrot, *Daucus carota*) to reveal gene duplicates originating from this whole genome duplication event (Supplemental Figure 2). Of the 32,431 genes that fell on chromosomes 1-8 of the *C. solstitialis* annotation, 8,184 (25.2%) were identified as putative paleologs (Supplemental Table 1). Single retained paleologs (genes inferred as arising during duplication, but for which other copies appear to be lost) were the most abundant class (4,911 genes) followed by paleologs retained in duplicate (3,036 genes) and triplicate (237 genes).

Pericentromeric regions can be predicted by areas of low gene density and high repeat content, particularly LTR-RTs (Scaglione et al. 2016). We used a sliding window analysis to assign relative scores in each window for low gene density or high repeat density, specifically based on unique repeat density, Gypsy LTR-RT density, and Copia LTR-RT density (Supplemental Figure 3). The average score of these categories revealed a single region of low gene density and high repeat density for each chromosome, indicating the location of the putative pericentromeric region (Figure 3). The sliding window analysis of gene density alone revealed roughly 2X higher gene density at the ends of chromosome arms relative to putative pericentromeric regions (Supplemental Figure 3).

**Figure 2.**
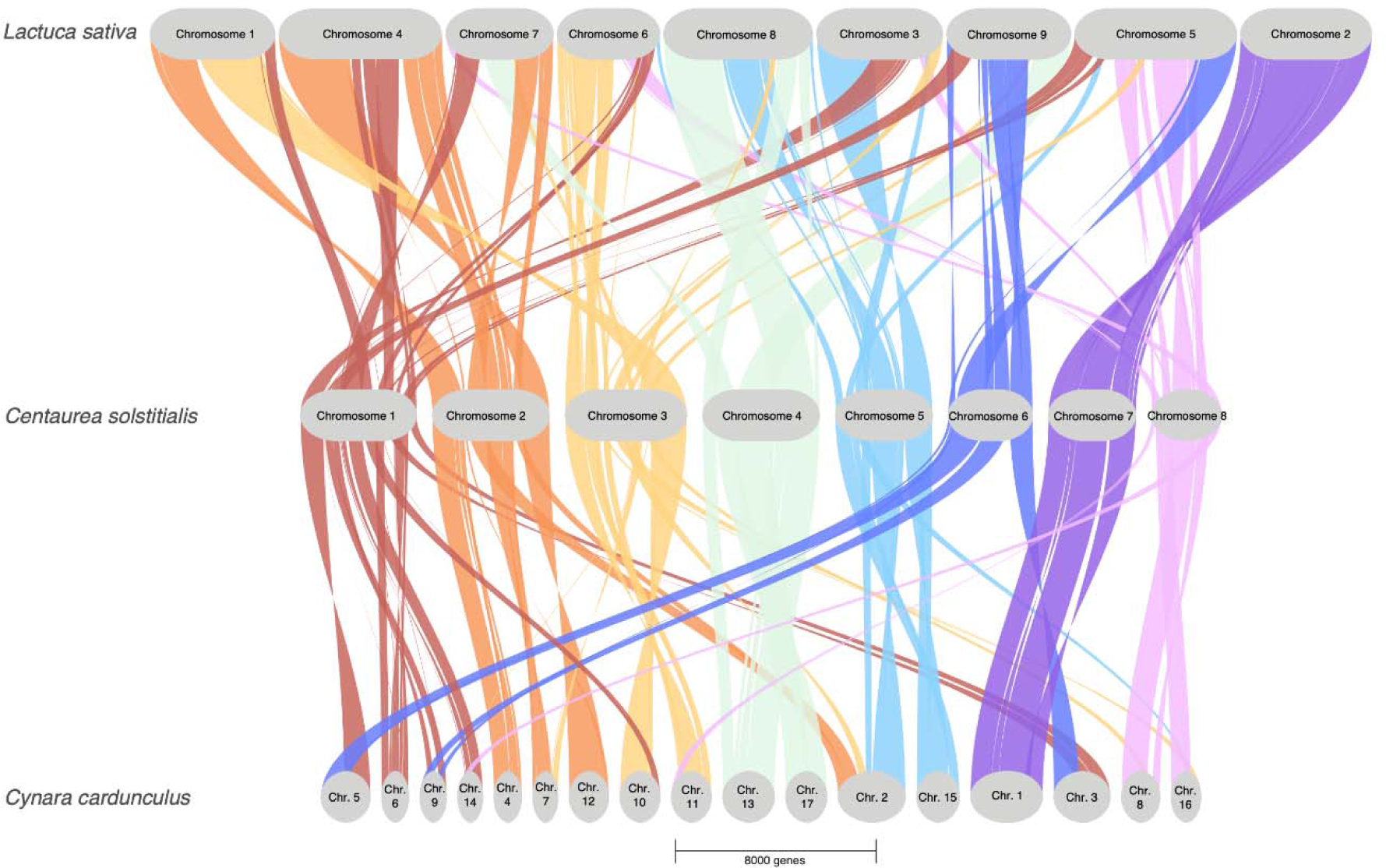
Gene synteny between yellow starthistle (*C. solstitialis*; 1N=8), globe artichoke (*Cynara cardunculus*; 1N=17), and lettuce (*Lactuca sativa*; 1N=9) genomes reveal chromosome evolution within the thistle subfamily (*Carduoideae*).

**Figure 3.**
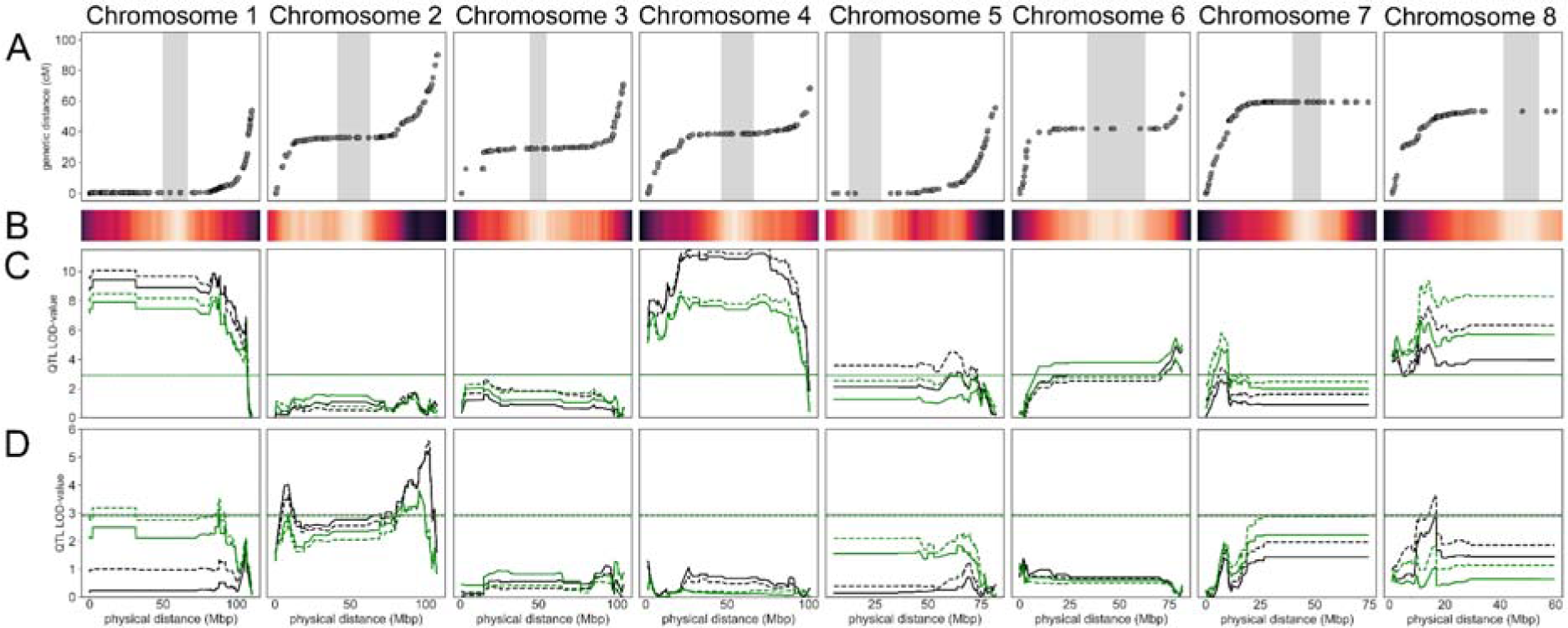
Summary of genome structure and QTL analyses. Panel A) Comparison of the physical map on the x-axis and genetic map on the y-axis reveals suppressed recombination in putative pericentromeric regions (vertical grey bars) for all chromosomes and putative structural variation on the arms of chromosomes 1 and 7. Panel B) centromere genome scan score heatmap of the average gene density, unique repetitive element density, Gypsy LTR-RT density, and Copia LTR-RT density across each chromosome. Panel C) QTL LOD scores for maximum leaf length at time point 1 (black lines) and time point 2 (green lines) using the HK method (dashed lines) and LOCO method (solid lines). Panel D) QTL LOD scores for total number of leaves at time point 1 (black lines) and time point 2 (green lines) using the HK method (dashed lines) and LOCO method (solid lines). Horizontal lines represent the suggestive QTL threshold of 0.1 for each time point and method.

Finally, the division of the genome into chromosomes can arise from fission, fusion, and duplication of chromosomal segments over evolutionary time, and this structure (and/or potential large scale errors in assembly) can be revealed by synteny comparisons with other closely related genomes (Wang et al. 2012). Gene synteny comparisons with globe artichoke (*Cynara cardunculus*; the most closely related thistle in subfamily *Carduoideae* whose genome has been assembled to the chromosome-scale) using GENESPACE revealed large blocks of conserved synteny across the genomes (Figure 2). Conserved synteny was also evident in comparisons between the *C. solstitialis* genome and the more divergent *Lactuca sativa* (subfamily *Cichorioideae*), albeit with more evidence for interchromosomal rearrangements (Figure 2). Three-way comparisons between *C. solstitialis*, *Cynara cardunculus,* and *Lactuca sativa* revealed a history of active chromosome evolution within the thistle subfamily *Carduoideae* (Figure 2), though *C. solstitialis* chromosome 7 was highly conserved across all three taxa (Figure 2).

### Genetic map

To validate the reference genome structure, characterize recombination patterns, and locate variable genetic markers across the genome, we constructed a genetic map using a population of 300 F2 individuals derived from a single cross between native and invading parents, genotyped using ddRADseq. RAD markers were aligned to the reference genome to identify variants, and then recombination rates among variants were used to infer a genetic map *de novo*, independent of the reference genome. After filtering for segregation distortion (removing markers based on a significance cutoff of p = 1E-5), a total of 1064 markers were variable and could be polarized as being either native or invader in origin. These formed eight linkage groups that corresponded with the eight chromosomes in the *C. solstitialis* reference genome (Figure 4B). Strong linkage and low estimated recombination were evident between markers within each chromosome, relative to linkage and recombination between markers on different chromosomes (Figure 4A). Only five aberrant markers from a single linkage group fell on a different reference assembly chromosome than the rest of the markers from their linkage group. These aberrant markers were pruned in the final map.

**Figure 4.**
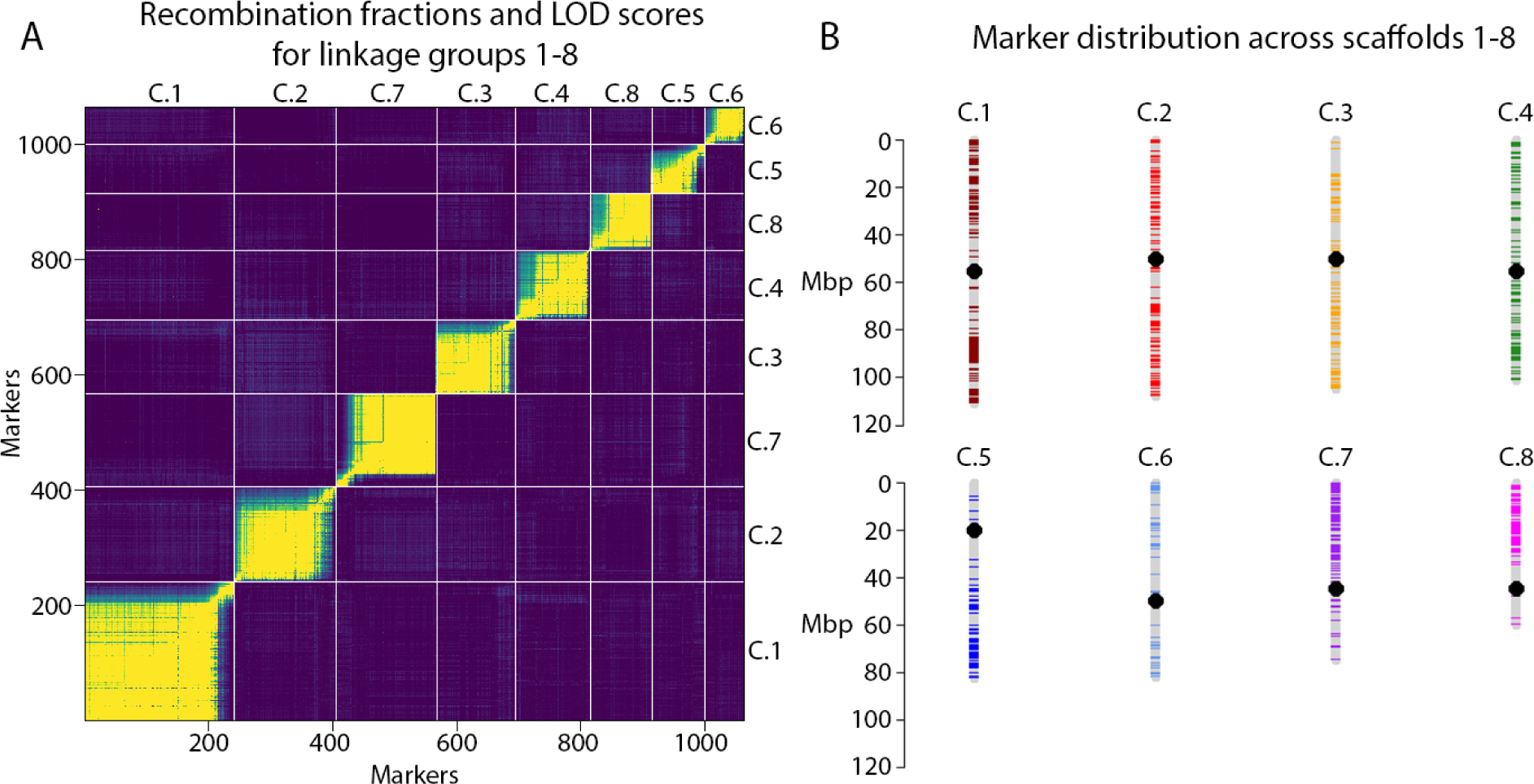
Visualization of the reference-aligned genetic map. A) Heatmap of recombination fractions in the top left of the diagonal and logarithm of odds (LOD) scores in the bottom right of the diagonal for ordered markers in linkage groups 1-8 reveal strong linkage and low recombination for markers within linkage groups and weak linkage with high recombination for markers on different linkage groups. Each linkage group is labeled with the corresponding scaffold (putative chromosome) number from the reference genome. B) Physical distribution of markers from the genetic map across chromosomes 1-8 of the reference genome, with each linkage group displayed as a different color. Putative centromere locations from genome scans are indicated with black dots.

Map distance comparisons between the genetic map and the reference assembly revealed megabase-scale regions of low recombination in the center of each chromosome, with chromosomes 1, 5, 7, and 8 also showing low recombination at one of the two ends of the chromosome (Figure 3A). For all chromosomes except 1 and 7, regions of low recombination in the genetic map corresponded closely with estimated pericentromeric regions (Figure 3A,B), and these same regions corresponded with lower RAD marker density (Figure 3A; Supplemental Figure 3). For chromosomes 1 and 7 recombination was low in putative pericentromeric regions but also on one of the distal arms of each chromosome (Figure 3A,B).

### Invader trait differences and QTL

We quantified growth differences between native and invading genotypes, and mapped QTL associated with these traits, using a common glasshouse experiment. This included 2901 F2 plants (including the 300 F2 plants used for the genetic map above), as well as 28-29 plants from each of the parental populations (P1), and an additional 41 F1s from bidirectional crosses between the parental populations. The plants were phenotyped for their total number of leaves and maximum leaf length at 3.5 and 5 weeks. For the P1 generation, invader genotypes had greater total number of leaves and greater maximum leaf length than native genotypes at both time points (Supplemental Figure 4), which is consistent with previous studies of invasive and native *C. solstitialis* populations (Dlugosch, Cang, et al. 2015). F1 individuals displayed heterosis for maximum leaf length at both time points, and for total number of leaves at five weeks (Supplemental Figure 4). F2 individuals showed the greatest range in trait values, as expected (Supplemental Figure 4). The 300 genotyped F2s were selected from the ends of the distribution of longest leaf length at 3.5 weeks, and they showed an expected bimodal distribution for this trait (Supplemental Figure 5). In contrast, the total number of leaves for those individuals followed a roughly normal distribution for both time points (Supplemental Figure 5).

Haley-Knott regression and leave one chromosome out (LOCO) genome scans revealed 13 distinct QTL peaks with varying degrees of support (summarized in Table 2). A total of seven peaks were associated with maximum leaf length only, four were associated with total number of leaves only, and two peaks were associated with both traits (i.e. putatively pleiotropic; Table 2). One of these two putatively pleiotropic QTL (peak 1 in Table 2) corresponded with the large region of low recombination outside of the pericentromeric region on chromosome 1 (Figure 3). Similarly, one of the suggestive QTL for total number of leaves (peak 11 in Table 2) corresponded with the other large region of low recombination outside of the pericentromeric region on chromosome 7 (Figure 3). These regions of low recombination - pericentromeric or otherwise - resulted in large QTL given that separate QTL were defined by a drop of at least 1 LOD from the peak. The number of candidate genes under QTL ranged from 3532 genes within the largest QTL spanning the 88.9 Mbp region of low recombination on chromosome 1, to 16 genes within the smallest QTL spanning 0.2 Mbp at the other end of chromosome 1 (Table 2, Supplemental Table 2).

**Table 2.** Summary of QTL peaks associated with maximum leaf length (len) and total number of leaves (num) at time points 1 and 2 (e.g. len1 and len2, respectively, or just len if the QTL is associated with the trait at both time points). The identification method, location in the reference, size of the peak, favored allele (i.e. TRI for invader, TK for native, and H for overdominance), percentage of variance explained (PVE), and number of candidate genes within 1 LOD drop of the peak are all given for each peak.

| PeakID | method | trait | chr | start | end | size (Mbp) | favored_allele | PVE | #candidates |
| --- | --- | --- | --- | --- | --- | --- | --- | --- | --- |
| 1 | LOCO | len_num | Chr_1 | 74748 | 88979819 | 88.90507 | H | 14.0 | 3532 |
| 2 | HK | len | Chr_1 | 106564625 | 106770853 | 0.206228 | H | 10.0 | 16 |
| 3 | LOCO | num | Chr_2 | 4353261 | 12526728 | 8.173467 | TRI | 6.0 | 495 |
| 4 | LOCO | num2 | Chr_2 | 80999070 | 99091674 | 18.0926 | TRI | 6.9 | 880 |
| 5 | LOCO | num1 | Chr_2 | 95334365 | 102637104 | 7.302739 | TRI | 7.7 | 370 |
| 6 | LOCO | len | Chr_4 | 1797495 | 7470976 | 5.673481 | TK | 11.6 | 407 |
| 7 | LOCO | len | Chr_4 | 20690837 | 75543191 | 54.85235 | TK | 15.7 | 2198 |
| 8 | LOCO | len1 | Chr_5 | 60237273 | 68394149 | 8.156876 | TK | 4.7 | 391 |
| 9 | LOCO | len | Chr_6 | 75934695 | 81496578 | 5.561883 | TK | 8.0 | 364 |
| 10 | LOCO | len | Chr_7 | 6148370 | 9845385 | 3.697015 | TRI | 7.0 | 282 |
| 11 | HK | num2 | Chr_7 | 19245630 | 74419950 | 55.17432 | TRI | 4.3 | 2102 |
| 12 | LOCO | len | Chr_8 | 1027923 | 2687378 | 1.659455 | TK | 8.2 | 147 |
| 13 | LOCO | len_num | Chr_8 | 10425376 | 16906147 | 6.480771 | TK | 9.6 | 353 |

Together, the 13 QTL peaks explained 85.9% and 82.5% of the phenotypic variation in maximum leaf length at 3.5 and 5 weeks, respectively, and 35.6% and 31.7% of the phenotypic variation in total number of leaves at 3.5 and 5 weeks, respectively. QTL peak 1 on chromosome 1 and peak 7 on chromosome 4 individually explained the most variation in maximum leaf length (14% and 15.7%, respectively), whereas peak 13 on chromosome 8 explained the most variation in total leaf number (9.6%; Table 2). For all four peaks associated with total number of leaves alone, the invader alleles at each QTL were associated with a greater number of leaves (Supplemental Figure 6). Of the seven peaks associated with maximum leaf length alone, the parental allele driving larger leaves varied between loci (Supplemental Figure 6). Overdominance was observed for both QTL peaks on chromosome 1 - including the peak that corresponded with the large region of low recombination - and the dominance and additive effects varied greatly among the other QTL (Supplemental Figure 6). Apparent deviations from additivity between QTL associated with the same trait indicated the possibility of epistatic interactions between some of the QTL (Supplemental Figure 7).

### Paleolog enrichment

Whole genome duplication results in syntenic blocks of duplicated genes that persist over evolutionary time, creating genomic regions of paleolog enrichment that might favor the generation of new QTL variants as a result of the evolution of one or more paleologs (Qi et al. 2021). Each QTL region will encompass multiple genes, and while we do not know the individual locus or loci responsible for the phenotypic effect of a QTL, we can test whether the genes under a given QTL are more likely to be paleologs than genes in other regions of the genome. To do this, we compared the frequency of paleologs within each QTL to genome-wide null distributions of randomly selected blocks of genes of the same size. From this analysis, two QTL peaks (peak 2 and peak 13) were found to be statistically enriched for paleologs (Supplemental Figure 8). Peak 2 contains only 16 genes, eight of which are paleologs. This includes Tryptophan aminotransferase-related protein 2 (TAR2), which is notable because it plays a role in auxin-dependent development processes such as growth in *Arabidopsis thaliana* (Stepanova et al. 2008), suggesting a strong potential to affect growth traits in *C. solstitialis*.

## Discussion

*Centaurea solstitialis* is a serious invasive pest of grasslands in the Americas, and our chromosome-scale assembly of its genome is the first for a non-cultivated species in the thistle subfamily (*Carduoideae*) of Asteraceae. We paired this reference genome with the first *C. solstitialis* QTL map to identify the genetic basis of evolving invader traits known from its well-studied ecology. Our results point to important roles of genome evolution during invasion, including potential impacts of overdominance, structural variants, and paleologs.

Assembly metrics, as well as an independently constructed genetic map, indicated that the eight primary scaffolds of our assembly constituted nearly complete pseudo-haploid chromosomes for *C. solstitialis*. These eight putative chromosomes contained 94.8% of the total sequence and 93.37% of complete BUSCO genes. The presence of only two additional BUSCO genes within the 1072 trailing scaffolds, and the fact that some BUSCO genes were instead duplicated within them, suggest that trailing scaffolds are likely largely composed of alternative haplotype or mis-assemblies due to heterozygosity in the outbred wild genotype used for genome assembly. Consistent with this hypothesis, only 16% of all annotated genes on the trailing scaffolds (294 genes total) were unique to those scaffolds and present in single copy - the rest had copies elsewhere in the genome. As a result, less than 1% of the total gene content of the genome was unique to the trailing scaffolds. Linkage information was also consistent with a chromosome-level assembly. The Omni-C contact map indicated that link density was higher within chromosomes than between, and that link density decayed with physical distance within a chromosome, as expected (Lajoie, Dekker, and Kaplan 2015). In addition, each of the eight chromosomes of the assembly corresponded with one of the eight linkage groups in the genetic map. Finally, the recombination frequencies between markers in the genetic map were consistent with the linear sequence of each chromosome in the assembly.

Annotations revealed a fairly typical genomic content, consistent with a relatively complete assembly. The chromosomes included 34,323 predicted gene models, comparable to other plant genomes, which have relatively little variation in gene number compared to their orders of magnitude variation in genome size (Wendel et al. 2016). Predictions included 616 tRNA models, similar to the 639 identified in *Arabidopsis thaliana* (Chan et al. 2021). Over 25% of genes were inferred to be paleologs derived from polyploidy at the base of the family, Asteraceae (M. S. Barker et al. 2016), which is a typical fraction of paleologs for angiosperms (Z. Li et al. 2021). Genome content is often dominated by repetitive DNA (Wendel et al. 2016), which was the case for *C. solstitialis*, with 63.3% of the genome occupied by repetitive elements. This included abundant long terminal repeat retrotransposons in the Ty1/Copia and Gypsy families. These results are similar to the repeat content of the globe artichoke genome which contains 58.4% repetitive DNA - with the most abundant families also being Ty1/Copia and Gypsy elements (Scaglione et al. 2016).

Importantly, recombination patterns in the mapping population revealed potential structural variants across genotypes. Chromosomes 1 and 7 displayed reduced recombination across one of their chromosome arms in the F2 population derived from an invader x native genotype cross, despite the central location of the inferred centromeres. Similar patterns have been explained by chromosomal rearrangements in a range of plant and animal taxa (Huang et al. 2020; Kirubakaran et al. 2016; C.-R. Lee et al. 2017; Tong et al. 2016), suggesting that native and invading genotypes of *C. solstitialis* might be characterized by large-scale structural variants on these chromosomes, relative to native genotypes. Notably, each of these regions of reduced recombination corresponded with one of the QTL peaks identified for morphological components of size variation, a trait that shows evidence for adaptive evolution and is thought to have aided in the invasion success of *C. solstitialis* (B. S. Barker et al. 2017; Dlugosch, Cang, et al. 2015). Structural variation is increasingly being recognized as an important source of genomic variation contributing to diverse ecological and evolutionary processes including adaptation and more recently invasion (Mérot et al. 2020; Huang and Rieseberg 2020; Battlay et al. 2022), and our results suggest it might play an important role in the rapid evolution of *C. solstitialis* invasions.

The region of reduced recombination on chromosome 1 was particularly intriguing because the corresponding QTL was both overdominant and putatively pleiotropic (i.e. associated with both maximum leaf length and total number of leaves). Given that heterosis is observed in the F1 generation for both traits, it is plausible that overdominance (and putative pleiotropy) at this QTL could be driven by many genes locked within a large non-recombining structural variant such as an inversion (Faria et al. 2019). Such a structural variant would prevent recombination across a large portion of the chromosome, which would “fix” heterozygous loci affecting both traits across its length in the F2 generation. All phenotypically relevant genes within this region would act as a single heterotic and pleiotropic locus. Indeed, one of the ways structural variation is known to contribute to adaptation is by creating strong linkage between adaptive alleles (Battlay et al. 2022), and balancing selection on overdominant inversions is a classic form of local adaptation in general (Faria et al. 2019). Alternatively, overdominance at this QTL could be explained by heterozygous loci across the inversion masking deleterious alleles (Connallon and Olito 2022; Jay et al. 2022), which are likely to accumulate during biological invasions due to strong genetic drift (Gilbert et al. 2017; Peischl, Kirkpatrick, and Excoffier 2015). In this case, the maintenance of the structural variant polymorphism within invasive populations would not need to be driven by local adaptation *per se*, but it could still be an important variant contributing to invasion success if invader genotypes that are heterozygous for the structural variant have higher fitness than homozygotes.

It is perhaps not surprising that QTL are associated with both of the regions of reduced recombination that we infer to be putative large-scale structural variants, given that any trait loci in these regions will be linked to all other loci across the chromosomal region. Even if a single large effect locus in proximity to the region was underpinning the association with size variation, the whole non-recombining region would still appear as a single QTL peak. Moreover, structural variants are predicted to have large phenotypic effects due to their potential to trap multiple tightly linked genes affecting a trait (Mérot et al. 2020). For this reason structural variants are predicted to be especially important components of adaptive variation during invasions because selection can favor these large effect regions despite the potential for strong founder effects and genetic drift to confound adaptation during range expansion (Bock et al. 2015; Dlugosch, Anderson, et al. 2015), or despite the potential for gene flow and admixture to homogenize local adaptation among founding populations (Reatini and Vision 2020). Population genomic analyses of linkage patterns within these chromosomes are needed to identify to what extent structural variants are present and limiting recombination within the invasion, and their potential contribution to the invasion success of *C. solstitialis*.

Gene duplications - whether single duplicated genes, blocks of duplicated genes, or whole genome duplications - are another important form of genome evolution that can contribute to adaptation and lead to evolutionary novelty (Conant and Wolfe 2008; Moriyama and Koshiba-Takeuchi 2018). Although mutations in duplicated genes can be masked initially due to functional redundancy, allowing them to accumulate, such mutations can generate either deleterious variation or novel adaptive variation that can experience selection in novel environments (Freeling 2009; Lynch and Force 2000; Baniaga et al. 2020). Interestingly, even gene duplicates that stem from relatively ancient whole genome duplication events occurring millions of years ago (paleologs) can contribute disproportionately to contemporary adaptive evolution - such as domestication alleles at loci originating from a ∼20 million year old whole genome duplication event in *Brassica rapa* (Qi et al. 2021). We found that two of our 13 QTL for invader traits in *C. solstitialis* were statistically enriched for paleologs, such that the specific gene or genes responsible for QTL phenotypic effects were disproportionately more likely to be paleologs than other genes in the genome. Of the two enriched QTL, one (peak 2) was associated with leaf length and displayed overdominance relative to parental genotypes. This peak is notable for being the smallest QTL we identified, consisting of only 16 genes, eight of which were paleologs. Among these paleologs was Tryptophan aminotransferase-related protein 2 (TAR2) which has been identified as playing a key role in auxin-dependent development processes in *Arabidopsis thaliana* - with mutants displaying phenotypic changes in both growth and reproduction (Stepanova et al. 2008). The other QTL that was statistically enriched for paleologs (peak 13) was also associated with both traits but the native genotype was generally larger than the invader genotype at this locus. Given that only two out of 13 QTL were enriched for paleologs, it is clear that paleolog variation alone is not responsible for rapid size evolution within the *C. solstitialis* invasion. Nevertheless, our results suggest that paleolog variation might supply important phenotypic variation under at least one invader QTL, and that the TAR2 ortholog should be queried further as a candidate gene for invader evolution.

Beyond the scope of invasion genomics, the *C. solstitialis* assembly is the second chromosome-scale reference genome for a member of the *Carduoideae*, which provides new opportunities to examine genome structural evolution in the Asteraceae. Gene synteny analyses among *C. solstitialis* and *Cynara cardunculus* (subfamily *Carduoideae*) and *Lactuca sativa* (subfamily *Cichorioideae*) revealed large regions of conserved synteny between all three genomes, as well as a history of dynamic rearrangements. The ancestral haploid chromosome number of *Asteraceae* is inferred to be 1N = 9 (Mota, Torices, and Loureiro 2016), which is retained in *Lactuca sativa* but not *C. solstitialis* (1N = 8) or *Cynara cardunculus* (1N = 17; Falistocco 2016). *Cynara cardunculus* has over double the chromosome number as *C. solstitialis*, yet Scaglione et al. (2016) found that no whole-genome duplication appears to have occurred after divergence between the *Cichorioideae* and *Carduoideae*, which occurred roughly 40 million years ago (Mandel et al. 2019). Consistent with Scaglione et al. (2016), our gene synteny analyses revealed evidence for widespread chromosome reorganization in *Cynara cardunculus* rather than whole-genome duplication. Karyotype analyses of *Cynara cardunculus* have revealed that the chromosomes display an unusual size distribution with discrete size categories of large, medium, and small chromosomes (Falistocco 2016), perhaps due to multiple fission events of larger ancestral chromosomes into smaller chromosomes. Our gene synteny analyses also suggest that the reduction in the chromosome number of *C. solstitialis* (1N = 8) relative to the base number for *Asteraceae* (1N = 9) may be due to many interchromosomal rearrangements between these genomes, underscoring the dynamic nature of genome evolution in the family.

Together, the reference genome assembly, annotation, genetic map, and QTL candidate regions presented here provide essential genomic resources for studies of *C. solstitialis* invasion, and for investigations of genome evolution across the agriculturally and evolutionarily important Asteraceae. Additional data could further develop the completeness of our reference and annotations, particularly additional whole genome sequence data to achieve a telomere-to- teleomore assembly and fill gaps, and pangenome sequencing of many individuals across the world wide distribution to characterize both nucleotide and structural variation (Li and Durbin 2024). Our findings here add to the growing appreciation of the potential for structural variants and gene duplicates to provide standing genetic variation for rapid adaptation. Future studies of population variation, patterns of linkage, and evolution at candidate loci can leverage this foundation to explore how different forms of genomic variation have contributed to the evolution of invasiveness in *Centaurea solstitialis* and expand to our understanding of rapid genome evolution more broadly.

## Materials and Methods

### Genomic reference sample collection and sequencing

Plant tissue used for the reference genome was grown from seed collected in August 2018 from the native range near Canales, Spain (site code ‘CAN’; Lat: 41.00033, Long: -4.89718). A reference voucher from the same locality is archived at the University of Arizona herbarium (ARIZ #425375). The plant was reared to the early bolting stage in a greenhouse at the University of Arizona (Tucson, Arizona, USA) under ambient light conditions from January to July 2020, and placed in the dark for 24 hours before harvesting leaves directly into liquid nitrogen for DNA extraction.

Leaf tissue was sent to Dovetail Genomics (Scotts Valley, California, USA) for reference genome sequencing and assembly using a combination of PacBio CLR and Dovetail Omni-C approaches. Specifically, PacBio CLR reads were generated by first constructing a ∼20kb library using SMRTbell Express Template Prep Kit 2.0 (PacBio, Menlo Park, CA, USA) using the manufacturer’s recommended protocol. The library was then bound to polymerase using the Sequel II Binding Kit 2.0 (PacBio) and sequenced on PacBio Sequel II 8M SMRT cells, generating 141 gigabases (Gb) of data, approximately 166x genome coverage. One Dovetail Omni-C library was prepared by fixing chromatin with formaldehyde, then digesting chromatin with DNAse I. Proximity ligation was then performed, crosslinks were reversed, and DNA was purified. The sequencing library was generated using NEBNext Ultra enzymes and Illumina-compatible adapters, followed by PCR enrichment. Libraries were sequenced on an Illumina HiSeqX platform, generating 59.8 Gb of data (approximately 71x genome coverage, or 35.5x per haplotype).

### Genome assembly

The initial assembly was constructed *de novo* from PacBio CLR reads using Wtdbg2 (Ruan and Li 2020). Potential contamination in the initial assembly was identified and removed based on BLAST v2.9 (Altschul et al. 1990) results against the NT database using Blobtools v1.1.1 (Laetsch and Blaxter 2017). The initial filtered assembly generated by Wtdbg2 was 1.06 Gb in length with a 742kb contig N50. Haplotypic duplications were then filtered from the assembly using purge_dups v1.1.2 (Guan et al. 2020), resulting in the removal of a total of 303Mb in 6099 contigs, for a final contig-level assembly of 764Mb with a 1.3Mb contig N50. The completeness of each phase of the assembly was evaluated using the 232655 Benchmarking Universal Single-Copy Orthologs (BUSCOs) from the Eudicot dataset (eudicot_odb10) from BUSCO (Manni et al. 2021) using compleasm v0.2.6 (Huang and Li, 2013). A total of 289 BUSCO genes that were identified as duplicated in the full Wtdbg2 assembly were single-copy in the purged assembly, and the purged assembly had a nearly identical total percentage of complete BUSCO genes (93.46% in the unpurged Wtdbg2 assembly, 92.99% in the purged assembly). The purged assembly was used as input along with the Dovetail Omni-C reads for scaffolding using the HiRise pipeline (Putnam et al. 2016). Briefly, Omni-C reads were aligned to the draft assembly, a likelihood model for genomic distance between read pairs was produced, and that model was then used to make joins and break misjoins in the input draft assembly. A total of 46.6 million valid read pairs were retained for scaffolding with HiRise, which introduced six contig breaks; the resulting assembly did not require misjoin adjustments.

### Genome annotation

Repeat family identification and classification were performed on the final genome assembly using RepeatModeler v2.0.1 (http://www.repeatmasker.org/RepeatModeler), relying on RECON v1.08 (Bao and Eddy 2002) and RepeatScout v1.0.6 (Price, Jones, and Pevzner 2005) for *de novo* identification of repeats. Custom repeat libraries produced by RepeatModeler were used to identify and mask repeats in the final assembly using RepeatMasker v4.1.0 (https://www.repeatmasker.org/RepeatMasker/). High-confidence transposable elements were then identified and classified by running RepeatMasker v4.1.0 with the plant TE database nrTEplantsApril2020 (Contreras-Moreira et al. 2021) as the input library. In brief, this library combines multiple plant TE databases and prunes the resulting library to remove redundant sequences and minimize overlap with protein-coding domains in nucleotide-binding, leucine-rich repeat (NLR) genes (Contreras-Moreira et al. 2021). Repetitive elements were then soft-masked in the reference.

For gene identification, transcripts from six *C. solstitialis* individuals were sequenced using RNAseq. Tissues included leaves from the same individual used for the reference genome, whole shoots of two additional small seedlings from the same population (‘CAN’), and three individuals (again including two seedlings and a mature bolting individual) from seed collected in September 2016 near Gilroy, California, USA in the invaded range (site code ‘GIL’; Lat: 37.03373, Long: -121.53674; reference voucher ARIZ #425113). All seeds were grown in early 2020 in greenhouses at the University of Arizona under ambient light conditions. Tissue was flash-frozen in liquid nitrogen and sent to Genewiz (Azenta Life Sciences, South Plainfield, NJew Jersey, USA) for RNA extraction and sequencing. RNA libraries were prepared via rRNA-depletion, and paired-end 2x150 reads were generated on the Illumina HiSeq platform.

Gene annotation was performed on the final repeat masked genome using a combination of AUGUSTUS v2.5.5 (Stanke et al. 2006) and SNAP v2006-07-28 (Korf 2004). Specifically, coding sequences from *Cynara cardunculus* (Acquadro et al. 2020), *Lactuca sativa* (Reyes-Chin-Wo et al. 2017), and *Helianthus annuus* (Badouin et al. 2017) were used to train two independent *ab initio* models for *C. solstitialis* using AUGUSTUS and SNAP. RNAseq data were then aligned to the final genome assembly using the STAR aligner software v2.7 (Dobin et al. 2013) and AUGUSTUS was used to generate intron hints using the *bam2hints* tool. Gene predictions were generated using MAKER (Holt and Yandell 2011), SNAP, and AUGUSTUS, using Swiss-Prot peptide sequences from the UniProt database to guide prediction and generate peptide evidence in the MAKER pipeline. The final set of genes was filtered to contain only genes predicted by both AUGUSTUS and SNAP. Putative gene function was assessed by performing a BLAST search of the peptide sequences against the UniProt database, and tRNA predictions were generated using tRNAscan-SE v2.05 (Chan et al. 2021). The quality of the annotation was evaluated by plotting annotation edit distance from MAKER2 and gene synteny was investigated between *C. solstitialis*, *Cynara cardunculus,* and *Lactuca sativa* using GENESPACE v1.3.1 (Lovell et al. 2022), which incorporates MCScanX (Wang et al. 2012) and OrthoFinder v2.5.4 (Emms and Kelly 2019). The genome assembly and annotation of globe artichoke were downloaded from http://www.artichokegenome.unito.it and the lettuce assembly (Lsat_Salinas_v11) and annotation were downloaded from NCBI (BioProject: PRJNA173551).

Sequence divergence among the genotypes used to construct the annotation was explored within gene regions to evaluate whether sequence divergence among the genotypes might impede alignment and annotation to the reference. RNAseq reads were re-aligned to the reference genome using the STAR aligner software v2.7 (Dobin et al. 2013) and SNPs were called from the resulting bam files using the mpileup function of bcftools v1.10.2. SNPs were filtered to only include those located within genes using bedtools v2.31.0, and then sequence differences between genotypes were identified using the isec function of bcftools for every pairwise comparison of the six genotypes used in the annotation. Total number of differences and per-base pair sequence divergence were calculated. In general, sequence divergence was low between all pairwise combinations of genotypes used to construct the annotation (Supplemental Table 3). Sequence divergence was on average higher among the invader genotypes than among native genotypes, and between native and invader genotypes (mean per-base pair divergence of 0.0061, 0.0048, 0.0057 and respectively) but there was also considerable variation in divergence among all of the genotypes (Supplemental Table 3). For instance, the western European reference genome genotype (CAN066) was more similar to one of the invader genotypes (GIL676) than either of the other western European genotypes (Supplemental Table 3).

### Paleolog identification

We identified blocks of paleologs using syntenic comparisons implemented in Frackify (McKibben and Barker 2021). Frackify utilizes CDS, GFF, and protein files from the *C. solstitialis* genome and an outgroup genome. *Daucus carota* (carrot; Dcarota_388_v2.0; Iorizzo et al. 2016) was selected as the outgroup because it is relatively closely related to *C. solstitialis* but does not share the known WGD event at the base of Asteraceae, which should permit the detection of paleologs stemming from that ancestral WGD in *C. solstitialis*. Additionally, *D. carot*a does not have its own WGD, which is preferred for inferring gene duplicates arising from WGD in the ingroup using Frackify (McKibben and Barker 2021). We used MCScanX to infer inter- and intraspecies syntenic blocks in the *C. solstitialis* and *D. carota* genomes (Wang et al. 2012). To identify ortholog divergences, a series of mixture models were fitted to the Ks distribution of interspecies collinear gene pairs using EMMIX (McLachlan and Peel 1999). The best fitting model based on the Bayesian Information Criterion determined the median Ks of ortholog divergences to be 1.40. Paralogs in the genome of *C. solstitialis* were identified using DupPipe and visualized in R using histograms (M. S. Barker et al. 2010). The median Ks of WGD peaks in the paralog age distribution was determined using EMMIX (McLachlan and Peel 1999). Syntenic inferences from MCScanX, orthology peak at Ks 1.40, and a WGD peak at median Ks 0.68 were used as inputs for Frackify.

### Identification of pericentromeric regions

Putative pericentromeric regions were identified using a sliding window analysis of gene and repeat density using a custom Python script‘*YST_genome.ipynb’*; available at (Reatini et al. 2022). Specifically, gene density from the gene annotation, unique repetitive element density from the repeat annotation, and Gypsy/Copia LTR-RT density from the curated TE database were all calculated within 5Mbp sized windows with a step size of 100,000 bp for each chromosome. The midpoint of each element was used to define its location in the genome. Counts for each of these four categories were normalized by dividing the count for the window by the maximum count for the category within the chromosome, yielding a measurement of the relative density of each category across the chromosome. Given that gene density is predicted to be low near centromeres but repeat density metrics are predicted to be high, the direction of the effect was rescaled such that higher values always corresponded with putative association with the centromere for all four categories. The average of the four categories was then calculated for each window in order to assign a score ranging from 0-1 to the window, with a score of 1 indicating the most centromere-associated scores across all four categories (minimum gene density, minimum unique repeat density, and maximum LTR-RT density for both Copia and Gypsy elements). This score was then plotted across each chromosome to estimate the location of pericentromeric regions.

### Mapping population collection and sequencing

An F2 mapping population was created using an initial cross between a single native range maternal parent and a single invaded range paternal parent. The native parent was grown from seed collected near Kirklareli, Turkey (site code ‘TK23’; Lat: 41.751233, Long: 27.247883) in September 2008. The invader parent was grown from seed collected near Mariposa, California, USA (site code ‘TRI’; Lat: 37.46178, Long: -119.79218) in 2008. Vouchers from each of these populations were deposited in the ARIZ herbarium (#425116-425117).

Parental plants were grown in a greenhouse at the University of British Columbia (Vancouver, British Columbia, Canada) and hand pollinated to produce F1 seed. Crosses were performed by covering flowering heads (capitula) with white organza bags before the heads opened to prevent pollination, then clipping heads presenting pollen and using these to brush against heads presenting a large fraction of receptive stigmas, and finally covering receiving heads again until seed maturation. Of the resulting F1 progeny, four were selected at random and reared to a large size in 11.4 L pots of commercial potting soil with 1 mL / L of 13:13:13 Osmocote fertilizer (Scotts Miracle-Gro, Marysville, Ohio, USA). The four F1 plants were reciprocally crossed to one another in two pairs, and samples of their leaves stored at -80 C for later genotyping.

Resulting F2 seeds were germinated on moist potting soil in an environmental room set to 12 hr days and 16-18 C days / 14 C nights, and misted by hand every other day. A total of 2901 F2 seedlings were transplanted to 410 ml Deepots (Steuwe & Sons, Tangent, Oregon, USA) in a 50:50 mix of silica sand and potting soil, with 2 mL of 13:13:13 Osmocote fertilizer. Deepots were watered daily from below on a flood table in a greenhouse at the University of British Columbia, under ambient lighting. In addition, 29 TK23 and 28 TRI plants from the parental populations, and 41 additional F1s from bidirectional crosses between them were included in the experiment for phenotyping. Of the 2901 F2 plants, 300 were selected for genotyping based on measurements of maximum leaf length at the first time point (see trait measurements below), including 150 plants with the longest maximum leaf length and 150 plants with the shortest maximum leaf length from the F2 generation.

Leaves from the 300 selected F2s, all four F1s, and six siblings of each parent (from field-collected seeds) were used for genotyping (parental tissue was not available for genotyping). DNA was extracted using a modified CTAB/PVP extraction protocol (Webb and Knapp 1990) and sent to Floragenex (Beaverton, Oregon, USA) for single digest Restriction Site Associated DNA (RAD) sequencing (Miller et al. 2007). RAD library preparation was performed with the CpNpG 5-methylcytosine sensitive enzyme, *PstI*, and 1x80 bp reads were sequenced on the Illumina Genome Analyzer II platform, yielding an average of 29.8X coverage per sample.

### Reference-guided genetic map

Raw single-end reads from the F2 mapping population were trimmed with the *process_radtags* function in STACKS v2.60 (Rochette, Rivera-Colón, and Catchen 2019) using the *c*, *q*, and *r* flags and specifying the *Pst*I restriction enzyme used in RAD library preparation. Trimmed reads were aligned to the largest eight scaffolds of the final reference assembly (reference chromosomes, see Results) using the *mem* function of the Burrows-Wheeler Aligner (BWA) (H. Li and Durbin 2009) and the resulting alignments were coordinate-sorted using samtools v1.10 (H. Li et al. 2009). RAD loci were built and genotyped using the reference-aligned pipeline of STACKS via the *ref_map.pl* wrapper. The *populations* function of STACKS was then used to filter the resulting genotype data, requiring that a locus be present in each generation (P1, F1, and F2) and with a minimum of 50% individuals represented at genotyped sites in order to be kept.

Filtered data were exported in variant call format, and parental genotypes for genetic mapping were called for each RAD marker using an allele frequency-based approach. A custom python script ‘*genmap_preprocessvcf.py’*; available at (Reatini et al. 2022) used the genotypes of six siblings of each parent to polarize the alleles present in F1s as derived from either the native or invading population, using the following criteria: for each allele present in the F1s at each RAD locus, the allele frequency among native (TK) sibs and invader (TRI) sibs was compared and if freq(TRI)>freq(TK) the allele was identified as predominantly invaded range (denoted ‘A’), whereas if freq(TK)>freq(TRI) the allele was identified as predominantly native range (denoted ‘B’). For a marker to be retained, all alleles in F1s needed to be successfully polarized, and there could be no more than two missing F1 genotypes or one missing parental population genotype. Filtered alleles were collapsed into parental classes (A and B), and F2 genotypes were exported to R/qtl format using these genotype calls. A total of 1,765 RAD markers were successfully polarized as being either native or invader in origin and used as input for initial linkage group formation in the genetic map.

Importantly, although the reference genome was used to call genotypes at each RAD marker, it was not used to group markers into linkage groups for genetic map construction. Instead, initial linkage groups for the genetic map were built *de novo* in R/qtl v1.50 (Broman et al. 2003) using the recombination fractions observed among F2 genotypes. Specifically, a maximum recombination fraction of 0.35 and minimum LOD score of 15 were used to group markers into initial linkage groups. Markers showing evidence of significant segregation distortion were removed using a significance cutoff of p=1E-5, yielding 1115 markers across 41 initial linkage groups. Of these markers, 1064 (95.4%) were located on the eight largest linkage groups, corresponding with the haploid chromosome number for *C. solstitialis*. Of the remaining linkage groups, 29 of them only held one marker each.

The eight largest linkage groups were used for the genetic map, assuming these linkage groups corresponded with the eight chromosomes of *C. solstitialis*. The genetic distances between markers in the genetic map were then estimated using R/qtl2 v0.28 (Broman et al. 2019, 2). Specifically, genetic distances were first estimated using the *est_map* function of R/qtl2 under a range of genotype error rates [0.001, 0.01, 0.025, 0.05, 0.075, 0.1]. The most likely error rate (0.075) was estimated using the log_10_ likelihood of each estimated map and this rate was used to estimate distances in the final genetic map.

To validate the linear sequence of the chromosomes in the reference genome, we plotted the collinearity between the genetic map and the physical reference using a custom Python script ‘*YST_genome.ipynb’*; available at Reatini et al. (2022). Aberrant markers in the genetic map - those that fell on different chromosomes than the rest of the linkage group - were quantified to determine the consistency between the reference genome and genetic map, and these aberrant markers were ultimately removed from downstream analyses. Collinearity patterns were then visually compared with centromere genome scans in order to evaluate whether patterns of recombination were consistent with the estimated centromere positions, again using custom Python scripts (*YST_genome.ipynb*; Reatini et al. 2022).

### Trait measurements and QTL analyses

The length of the longest leaf and the total number of leaves were recorded for all 2901 plants at 3.5 weeks and 5 weeks of age. Potential confounding factors during trait measurement were accounted for by fitting linear models for each trait measurement using the *aov* function in R, with fixed effects of the day measurements were collected, the individual collecting the data, the spatial location in the greenhouse (block), the location of the plant within the block, and the identity of the specific F1 mother of each F2. Residuals of the model were then extracted and added to the grand mean to obtain corrected values of maximum leaf length and total number of leaves at both time points for use in QTL analysis.

Additional representatives of both parental populations and F1 crosses between these populations were included in the experiment for phenotypic comparison to the F2s. Parental population plants were grown from field-collected seed including 28 ‘TRI’ plants from the invasion and 29 ‘TK23’ plants from the native range. F1 seeds came from additional controlled crosses, and included 17 plants with ‘TRI’ genotypes as the maternal parents and 24 plants with ‘TK23’ genotypes as the maternal parents. Germination and rearing were concurrent with the F2 populations, under the same conditions.

Genome scans to identify QTL were performed using both Haley-Knott regression (Haley and Knott 1992) and the leave one chromosome out (Yang et al. 2014) approach in R/qtl2 using the final genetic map and corrected phenotypic data as input. Both models were used to qualitatively gauge support for QTL by comparing overlap between them. Given that the LOCO method accounts for kinship when crosses are generated with parents from divergent source populations – as was the case for the present mapping population – the LOCO model was used as the primary model for candidate gene identification. For each method, two thresholds were used to define QTL peaks using a permutation-based approach in R/qtl2: a standard significance threshold of 0.05 and a suggestive threshold of 0.1. Separate peaks on the same chromosome were identified using a minimum LOD decline of 1 between peaks. Candidate genes within QTL peaks were identified by extracting all genes within a decline of 1 LOD on either side of the peak.

To quantify the phenotypic effects of each QTL peak identified in the above scans, the percentage of variance explained was calculated using the formula PVE == 1 - 10^-(2/n)LOD^ where *n* was the sample size of 300 F2 individuals from the mapping population following Broman and Sen (2009). Genotypic, additive, and dominance effects were then estimated using the *scan1coef* function of R/qtl2 to quantify the direction and magnitude of each effect. Epistatic interactions between QTL associated with the same traits were then explored by first extracting genotypes for each QTL peak across all F2s using the *maxmarg* function of R/qtl2. For each pairwise combination of QTL associated with the same trait, the mean phenotype was then plotted for each two-locus combination of genotypes in order to qualitatively assess deviations from additivity.

### Paleolog enrichment analysis

To assess whether any QTL were enriched for paleologs, null distributions of paleologs across the genome were built by assessing the frequency of paleologs within randomly selected blocks of genes the size of each QTL. Specifically, for each QTL, the number of genes within the QTL was recorded, and then 100,000 blocks of consecutive genes of the same size as the QTL were randomly sampled from across the genome, with the frequency of paleologs calculated within each block. To assess whether the QTL was in the tail of the null distribution, the frequency of paleologs within the QTL was z-tranformed using the mean and standard deviation from the null distribution. Two-tailed p-values were then calculated from these Z-scores to assess significance, and the distributions for each QTL were plotted in R for visualization.

## Acknowledgements

We thank B.M. Anderson, S. Lin and K. Nurkowski for help with data collection and D. Kaplan for assistance with greenhouse logistics. This work was supported by funding from a Natural Sciences and Engineering Research Council of Canada grant #353026 to L.H.R., United States National Science Foundation grants #1750280 to KMD and #2109625 to BR, and United States Department of Agriculture grant #2023-67013-40169 to KMD.

## Data Availability

The chromosome-scale genome assembly,annotation, and all raw sequencing data used to construct them have been deposited at GenBank under BioProject PRJNA902738 (assembly ASM3016916v1). All custom analysis code and all results necessary to replicate the analyses and plots presented here are available at zenodo - [restricted access for peer review: doi: https://doi.org/10.5281/zenodo.7324093].

## Author Contributions

BR, JP, KD, MB, and LR conceived the work. BR, JP, KD, AC, QJ, and MM conducted the research and analyzed the resulting data. BR and KD drafted the manuscript, and all authors provided revisions and feedback.

## References

Acquadro, Alberto, Ezio Portis, Danila Valentino, Lorenzo Barchi, and Sergio Lanteri. 2020. “‘Mind the Gap’: Hi-C Technology Boosts Contiguity of the Globe Artichoke Genome in Low-Recombination Regions.” G3: Genes|Genomes|Genetics 10 (10): 3557–64. 10.1534/g3.120.401446.

Altschul, S. F., Gish, W., Miller, W., Myers, E. W., & Lipman, D. J. 1990. Basic local alignment search tool. Journal of Molecular Biology 215 (3): 403–410.

Badouin, Hélène, Jérôme Gouzy, Christopher J. Grassa, Florent Murat, S. Evan Staton, Ludovic Cottret, Christine Lelandais-Brière, et al. 2017. “The Sunflower Genome Provides Insights into Oil Metabolism, Flowering and Asterid Evolution.” Nature 546 (7656): 148–52. 10.1038/nature22380.

Bancheva, S., and J. Greilhuber. 2006. “Genome Size in Bulgarian Centaurea s.l. (Asteraceae).” Plant Systematics and Evolution 257 (1): 95–117. 10.1007/s00606-005-0384-7.

Baniaga, Anthony E., Hannah E. Marx, Nils Arrigo, and Michael S. Barker. 2020. “Polyploid Plants Have Faster Rates of Multivariate Niche Differentiation than Their Diploid Relatives.” Ecology Letters 23 (1): 68–78. 10.1111/ele.13402.

Bao, Zhirong, and Sean R. Eddy. 2002. “Automated De Novo Identification of Repeat Sequence Families in Sequenced Genomes.” Genome Research 12 (8): 1269–76. 10.1101/gr.88502.

Barker, Brittany S., Krikor Andonian, Sarah M. Swope, Douglas G. Luster, and Katrina M. Dlugosch. 2017. “Population Genomic Analyses Reveal a History of Range Expansion and Trait Evolution across the Native and Invaded Range of Yellow Starthistle (Centaurea Solstitialis).” Molecular Ecology 26 (4): 1131–47. 10.1111/mec.13998.

Barker, Brittany S., Janelle E. Cocio, Samantha R. Anderson, Joseph E. Braasch, Feng A. Cang, Heather D. Gillette, and Katrina M. Dlugosch. 2019. “Potential Limits to the Benefits of Admixture during Biological Invasion.” Molecular Ecology 28 (1): 100–113. 10.1111/mec.14958.

Barker, Michael S., Katrina M. Dlugosch, Louie Dinh, R. Sashikiran Challa, Nolan C. Kane, Matthew G. King, and Loren H. Rieseberg. 2010. “EvoPipes.Net: Bioinformatic Tools for Ecological and Evolutionary Genomics.” Evolutionary Bioinformatics Online 6 (October): 143–49. 10.4137/EBO.S5861.

Barker, Michael S., Nolan C. Kane, Marta Matvienko, Alexander Kozik, Richard W. Michelmore, Steven J. Knapp, and Loren H. Rieseberg. 2008. “Multiple Paleopolyploidizations during the Evolution of the Compositae Reveal Parallel Patterns of Duplicate Gene Retention after Millions of Years.” Molecular Biology and Evolution 25 (11): 2445–55. 10.1093/molbev/msn187.

Barker, Michael S., Zheng Li, Thomas I. Kidder, Chris R. Reardon, Zhao Lai, Luiz O. Oliveira, Moira Scascitelli, and Loren H. Rieseberg. 2016. “Most Compositae (Asteraceae) Are Descendants of a Paleohexaploid and All Share a Paleotetraploid Ancestor with the Calyceraceae.” American Journal of Botany 103 (7): 1203–11. 10.3732/ajb.1600113.

Battlay, Paul, Jonathan Wilson, Vanessa C. Bieker, Chris Lee, Diana Prapas, Bent Petersen, Sam Craig, et al. 2022. “Large Haploblocks Underlie Rapid Adaptation in an Invasive Weed.” bioRxiv. 10.1101/2022.03.02.482376.

Beest, Mariska te, Johannes J. Le Roux, David M. Richardson, Anne K. Brysting, Jan Suda, Magdalena Kubešová, and Petr Pyšek. 2012. “The More the Better? The Role of Polyploidy in Facilitating Plant Invasions.” Annals of Botany 109 (1): 19–45. 10.1093/aob/mcr277.

Bock, Dan G., Celine Caseys, Roger D. Cousens, Min A. Hahn, Sylvia M. Heredia, Sariel Hübner, Kathryn G. Turner, Kenneth D. Whitney, and Loren H. Rieseberg. 2015. “What We Still Don’t Know about Invasion Genetics.” Molecular Ecology 24 (9): 2277–97. 10.1111/mec.13032.

Broman, Karl W, Daniel M Gatti, Petr Simecek, Nicholas A Furlotte, Pjotr Prins, Śaunak Sen, Brian S Yandell, and Gary A Churchill. 2019. “R/Qtl2: Software for Mapping Quantitative Trait Loci with High-Dimensional Data and Multiparent Populations.” Genetics 211 (2): 495–502. 10.1534/genetics.118.301595.

Broman, Karl W., and Saunak Sen. 2009. A Guide to QTL Mapping with R/Qtl. Statistics for Biology and Health. Dordrecht: Springer.

Broman, Karl W., Hao Wu, Śaunak Sen, and Gary A. Churchill. 2003. “R/Qtl: QTL Mapping in Experimental Crosses.” Bioinformatics 19 (7): 889–90. 10.1093/bioinformatics/btg112.

Cang, F. A., Welles, S. R., Wong, J., Ziaee, M., and Dlugosch, Katrina M. 2024. “Genome size variation and evolution during invasive range expansion in an introduced plant”. Evolutionary Applications 17, e13624. 10.1111/eva.13624.

Chan, Patricia P, Brian Y Lin, Allysia J Mak, and Todd M Lowe. 2021. “tRNAscan-SE 2.0: Improved Detection and Functional Classification of Transfer RNA Genes.” Nucleic Acids Research 49 (16): 9077–96. 10.1093/nar/gkab688.

Conant, Gavin C., and Kenneth H. Wolfe. 2008. “Turning a Hobby into a Job: How Duplicated Genes Find New Functions.” Nature Reviews Genetics 9 (12): 938–50. 10.1038/nrg2482.

Connallon, Tim, and Colin Olito. 2022. “Natural Selection and the Distribution of Chromosomal Inversion Lengths.” Molecular Ecology 31 (13): 3627–41. 10.1111/mec.16091.

Contreras-Moreira, Bruno, Carla V Filippi, Guy Naamati, Carlos García Girón, James E Allen, and Paul Flicek. 2021. “K-Mer Counting and Curated Libraries Drive Efficient Annotation of Repeats in Plant Genomes.” The Plant Genome 14 (3): e20143. 10.1002/tpg2.20143.

Dlugosch, Katrina M., Samantha R. Anderson, Joseph Braasch, F. Alice Cang, and Heather D. Gillette. 2015. “The Devil Is in the Details: Genetic Variation in Introduced Populations and Its Contributions to Invasion.” Molecular Ecology 24 (9): 2095–2111. 10.1111/mec.13183.

Dlugosch, Katrina M., F. Alice Cang, Brittany S. Barker, Krikor Andonian, Sarah M. Swope, and Loren H. Rieseberg. 2015. “Evolution of Invasiveness through Increased Resource Use in a Vacant Niche.” Nature Plants 1 (June). 10.1038/nplants.2015.66.

Dobin, Alexander, Carrie A. Davis, Felix Schlesinger, Jorg Drenkow, Chris Zaleski, Sonali Jha, Philippe Batut, Mark Chaisson, and Thomas R. Gingeras. 2013. “STAR: Ultrafast Universal RNA-Seq Aligner.” Bioinformatics 29 (1): 15–21. 10.1093/bioinformatics/bts635.

Emms, David M., and Steven Kelly. 2019. “OrthoFinder: phylogenetic orthology inference for comparative genomics.” Genome Biology 20: 238.

Eriksen, Renée L., Theodora Desronvil, José L. Hierro, and Rick Kesseli. 2012. “Morphological Differentiation in a Common Garden Experiment among Native and Non-Native Specimens of the Invasive Weed Yellow Starthistle (Centaurea Solstitialis).” Biological Invasions 14 (7): 1459–67. 10.1007/s10530-012-0172-6.

Estoup, Arnaud, Virginie Ravigné, Ruth Hufbauer, Renaud Vitalis, Mathieu Gautier, and Benoit Facon. 2016. “Is There a Genetic Paradox of Biological Invasion?” Annual Review of Ecology, Evolution, and Systematics 47 (1): 51–72. 10.1146/annurev-ecolsys-121415-032116.

Falistocco, Egizia. 2016. “Cytogenetic Characterization of Cultivated Globe Artichoke (Cynara Cardunculus Var. Scolymus) and Cardoon (C. Cardunculus Var. Altilis).” Caryologia 69 (1): 1–4. 10.1080/00087114.2015.1109935.

Faria, Rui, Kerstin Johannesson, Roger K. Butlin, and Anja M. Westram. 2019. “Evolving Inversions.” Trends in Ecology & Evolution 34 (3): 239–48. 10.1016/j.tree.2018.12.005.

Freeling, Michael. 2009. “Bias in Plant Gene Content Following Different Sorts of Duplication: Tandem, Whole-Genome, Segmental, or by Transposition.” Annual Review of Plant Biology 60 (1): 433–53. 10.1146/annurev.arplant.043008.092122.

Gerlach, John D. 1997. “How the West Was Lost: Reconstructing the Invasion Dynamics of Yellow Starthistle and Other Plant Invaders of Western Rangelands and Natural Areas.” Calif. Exot. Pest Plant Counc. Symp. Proc., 6.

Gilbert, Kimberly J., Nathaniel P. Sharp, Amy L. Angert, Gina L. Conte, Jeremy A. Draghi, Frédéric Guillaume, Anna L. Hargreaves, Remi Matthey-Doret, and Michael C. Whitlock. 2017. “Local Adaptation Interacts with Expansion Load during Range Expansion: Maladaptation Reduces Expansion Load.” The American Naturalist 189 (4): 368–80. 10.1086/690673.

Guan, Dengfeng, Shane A McCarthy, Jonathan Wood, Kerstin Howe, Yadong Wang, and Richard Durbin. 2020. “Identifying and Removing Haplotypic Duplication in Primary Genome Assemblies.” Bioinformatics 36 (9): 2896–98. 10.1093/bioinformatics/btaa025.

Haley, C. S., and S. A. Knott. 1992. “A Simple Regression Method for Mapping Quantitative Trait Loci in Line Crosses Using Flanking Markers.” Heredity 69 (4): 315–24. 10.1038/hdy.1992.131.

Heiser Jr., Charles B., and Thomas W. Whitaker. 1948. “Chromosome Number, Polyploidy, and Growth Habit in California Weeds.” American Journal of Botany 35 (3): 179–86. 10.1002/j.1537-2197.1948.tb05204.x.

Holt, Carson, and Mark Yandell. 2011. “MAKER2: An Annotation Pipeline and Genome-Database Management Tool for Second-Generation Genome Projects.” BMC Bioinformatics 12 (1): 491. 10.1186/1471-2105-12-491.

Huang, Kaichi, Rose L. Andrew, Gregory L. Owens, Kate L. Ostevik, and Loren H. Rieseberg. 2020. “Multiple Chromosomal Inversions Contribute to Adaptive Divergence of a Dune Sunflower Ecotype.” Molecular Ecology 29 (14): 2535–49. 10.1111/mec.15428.

Huang, Kaichi, and Loren H. Rieseberg. 2020. “Frequency, Origins, and Evolutionary Role of Chromosomal Inversions in Plants.” Frontiers in Plant Science 11. https://www.frontiersin.org/articles/10.3389/fpls.2020.00296.

Huang, Neng, and Heng Li. 2023. “compleasm: a faster and more accurate reimplementation of BUSCO.” Bioinformatics 39: btad595. doi:10.1093/bioinformatics/btad595.

Iorizzo, Massimo, Shelby Ellison, Douglas Senalik, Peng Zeng, Pimchanok Satapoomin, Jiaying Huang, Megan Bowman, et al. 2016. “A High-Quality Carrot Genome Assembly Provides New Insights into Carotenoid Accumulation and Asterid Genome Evolution.” Nature Genetics 48 (6): 657–66. 10.1038/ng.3565.

Irimia, Ramona-Elena, Daniel Montesinos, Özkan Eren, Christopher J. Lortie, Kristine French, Lohengrin A. Cavieres, Gastón J. Sotes, José L. Hierro, Andreia Jorge, and João Loureiro. 2017. “Extensive Analysis of Native and Non-Native Centaurea Solstitialis L. Populations across the World Shows No Traces of Polyploidization.” PeerJ 5 (August): e3531. 10.7717/peerj.3531.

Jay, Paul, Emilie Tezenas, Amandine Véber, and Tatiana Giraud. 2022. “Sheltering of Deleterious Mutations Explains the Stepwise Extension of Recombination Suppression on Sex Chromosomes and Other Supergenes.” PLOS Biology 20 (7): e3001698. 10.1371/journal.pbio.3001698.

Kirubakaran, Tina Graceline, Harald Grove, Matthew P. Kent, Simen R. Sandve, Matthew Baranski, Torfinn Nome, Maria Cristina De Rosa, et al. 2016. “Two Adjacent Inversions Maintain Genomic Differentiation between Migratory and Stationary Ecotypes of Atlantic Cod.” Molecular Ecology 25 (10): 2130–43. 10.1111/mec.13592.

Korf, Ian. 2004. “Gene Finding in Novel Genomes.” BMC Bioinformatics 5 (1): 59. 10.1186/1471-2105-5-59.

Laetsch, Dominik R., and Mark L. Blaxter. 2017. “BlobTools: Interrogation of Genome Assemblies.” F1000Research. 10.12688/f1000research.12232.1.

Lajoie, Bryan R., Job Dekker, and Noam Kaplan. 2015. “The Hitchhiker’s Guide to Hi-C Analysis: Practical Guidelines.” *Methods (San Diego*, Calif.) 72 (January): 65–75. 10.1016/j.ymeth.2014.10.031.

Lee, Carol Eunmi. 2002. “Evolutionary Genetics of Invasive Species.” Trends in Ecology & Evolution 17 (8): 386–91. 10.1016/S0169-5347(02)02554-5.

Lee, Cheng-Ruei, Baosheng Wang, Julius P. Mojica, Terezie Mandáková, Kasavajhala V. S. K. Prasad, Jose Luis Goicoechea, Nadeesha Perera, et al. 2017. “Young Inversion with Multiple Linked QTLs under Selection in a Hybrid Zone.” Nature Ecology & Evolution 1 (5): 1–13. 10.1038/s41559-017-0119.

Li, Heng, and Richard Durbin. 2009. “Fast and Accurate Short Read Alignment with Burrows–Wheeler Transform.” Bioinformatics 25 (14): 1754–60. 10.1093/bioinformatics/btp324.

Li, Heng, and Richard Durbin. 2024. “Genome assembly in the telomere-to-telomere era”. Nat. Rev. Genet. 25, 658–670. 10.1038/s41576-024-00718-w.

Li, Heng, Bob Handsaker, Alec Wysoker, Tim Fennell, Jue Ruan, Nils Homer, Gabor Marth, Goncalo Abecasis, and Richard Durbin. 2009. “The Sequence Alignment/Map Format and SAMtools.” Bioinformatics 25 (16): 2078–79. 10.1093/bioinformatics/btp352.

Li, Zheng, Michael T.W. McKibben, Geoffrey S. Finch, Paul D. Blischak, Brittany L. Sutherland, and Michael S. Barker. 2021. “Patterns and Processes of Diploidization in Land Plants.” Annual Review of Plant Biology 72 (1): 387–410. 10.1146/annurev-arplant-050718-100344.

Lovell, John T., Avinash Sreedasyam, M. Eric Schranz, Melissa Wilson, Joseph W Carlson, Alex Harkess, David Emms, David M. Goodstein, and Jeremy Schmutz. 2022. “GENESPACE tracks regions of interest and gene copy number variation across multiple genomes.” eLife 11:e78526.

Lynch, Michael, and Allan G. Force. 2000. “The Origin of Interspecific Genomic Incompatibility via Gene Duplication.” The American Naturalist 156 (6): 590–605. 10.1086/316992.

Maddox, Donald M., Aubrey Mayfield, and Noah H. Poritz. 1985. “Distribution of Yellow Starthistle (Centaurea Solstitialis) and Russian Knapweed (Centaurea Repens).” Weed Science 33 (3): 315– 27.

Mandel, Jennifer R., Rebecca B. Dikow, Carolina M. Siniscalchi, Ramhari Thapa, Linda E. Watson, and Vicki A. Funk. 2019. “A Fully Resolved Backbone Phylogeny Reveals Numerous Dispersals and Explosive Diversifications throughout the History of Asteraceae.” Proceedings of the National Academy of Sciences 116 (28): 14083–88. 10.1073/pnas.1903871116.

Manni, Mosè, Matthew R Berkeley, Mathieu Seppey, Felipe A Simão, and Evgeny M Zdobnov. 2021. “BUSCO Update: Novel and Streamlined Workflows along with Broader and Deeper Phylogenetic Coverage for Scoring of Eukaryotic, Prokaryotic, and Viral Genomes.” Molecular Biology and Evolution 38 (10): 4647–54. 10.1093/molbev/msab199.

McGaughran, Angela, Manpreet K Dhami, Elahe Parvizi, Amy L Vaughan, Dianne M Gleeson, Kathryn A Hodgins, Lee A Rollins, et al. 2024. “Genomic Tools in Biological Invasions: Current State and Future Frontiers.” Genome Biology and Evolution 16 (1): evad230. 10.1093/gbe/evad230.

McKibben, Michael T. W., and Michael S. Barker. 2021. “Applying Machine Learning to Classify the Origins of Gene Duplications.” bioRxiv. 10.1101/2021.08.12.456144.

McLachlan, Geoff, and David Peel. 1999. “The EMMIX Algorithm for the Fitting of Normal and T-Components.” Journal of Statistical Software 4 (July): 1–14. 10.18637/jss.v004.i02.

Mérot, Claire, Rebekah A. Oomen, Anna Tigano, and Maren Wellenreuther. 2020. “A Roadmap for Understanding the Evolutionary Significance of Structural Genomic Variation.” Trends in Ecology & Evolution 35 (7): 561–72. 10.1016/j.tree.2020.03.002.

Miller, M. R., J. P. Dunham, A. Amores, W. A. Cresko, and E. A. Johnson. 2007. “Rapid and Cost-Effective Polymorphism Identification and Genotyping Using Restriction Site Associated DNA (RAD) Markers.” Genome Research 17 (2): 240–48. 10.1101/gr.5681207.

Montesinos, Daniel, and Ragan M. Callaway. 2017. “Inter-Regional Hybrids of Native and Invasive Centaurea Solstitialis Display Intermediate Competitive Ability.” Ecography 40 (7): 801–2. 10.1111/ecog.02653.

Montesinos, Daniel, and Ragan M. Callaway. 2018. “Traits Correlate with Invasive Success More than Plasticity: A Comparison of Three Centaurea Congeners.” Ecology and Evolution 8 (15): 7378–85. 10.1002/ece3.4080.

Montesinos, Daniel, Ryan C. Graebner, and Ragan M. Callaway. 2019. “Evidence for Evolution of Increased Competitive Ability for Invasive Centaurea Solstitialis, but Not for Naturalized C. Calcitrapa.” Biological Invasions 21 (1): 99–110. 10.1007/s10530-018-1807-z.

Moriyama, Yuuta, and Kazuko Koshiba-Takeuchi. 2018. “Significance of Whole-Genome Duplications on the Emergence of Evolutionary Novelties.” Briefings in Functional Genomics 17 (5): 329–38. 10.1093/bfgp/ely007.

Mota, Lucie, Rubén Torices, and João Loureiro. 2016. “The Evolution of Haploid Chromosome Numbers in the Sunflower Family.” Genome Biology and Evolution 8 (11): 3516–28. 10.1093/gbe/evw251.

Mounger, Jeannie, Malika L. Ainouche, Oliver Bossdorf, Armand Cavé-Radet, Bo Li, Madalin Parepa, Armel Salmon, Ji Yang, and Christina L. Richards. 2021. “Epigenetics and the Success of Invasive Plants.” Philosophical Transactions of the Royal Society B: Biological Sciences 376 (1826): 20200117. 10.1098/rstb.2020.0117.

Nei, Masatoshi, Takeo Maruyama, and Ranajit Chakraborty. 1975. “The Bottleneck Effect and Genetic Variability in Populations.” Evolution 29 (1): 1–10. 10.2307/2407137.

Niu, Xiao-Min, Yong-Chao Xu, Zi-Wen Li, Yu-Tao Bian, Xing-Hui Hou, Jia-Fu Chen, Yu-Pan Zou, et al. 2019. “Transposable Elements Drive Rapid Phenotypic Variation in Capsella Rubella.” Proceedings of the National Academy of Sciences 116 (14): 6908–13. 10.1073/pnas.1811498116.

Orr, H. Allen. 1998. “The Population Genetics of Adaptation: The Distribution of Factors Fixed during Adaptive Evolution.” Evolution 52 (4): 935. 10.2307/2411226.

Peischl, Stephan, Isabelle Dupanloup, Adrien Foucal, Michèle Jomphe, Vanessa Bruat, Jean-Christophe Grenier, Alexandre Gouy, et al. 2018. “Relaxed Selection During a Recent Human Expansion.” Genetics 208 (2): 763–77. 10.1534/genetics.117.300551.

Peischl, Stephan, Mark Kirkpatrick, and Laurent Excoffier. 2015. “Expansion Load and the Evolutionary Dynamics of a Species Range.” The American Naturalist 185 (4): E81–93. 10.1086/680220.

Price, Alkes L., Neil C. Jones, and Pavel A. Pevzner. 2005. “De Novo Identification of Repeat Families in Large Genomes.” *Bioinformatics (Oxford,* England) 21 Suppl 1 (June): i351-358. 10.1093/bioinformatics/bti1018.

Putnam, Nicholas H., Brendan L. O’Connell, Jonathan C. Stites, Brandon J. Rice, Marco Blanchette, Robert Calef, Christopher J. Troll, et al. 2016. “Chromosome-Scale Shotgun Assembly Using an in Vitro Method for Long-Range Linkage.” Genome Research 26 (3): 342–50. 10.1101/gr.193474.115.

Qi, Xinshuai, Hong An, Tara E. Hall, Chenlu Di, Paul D. Blischak, Michael T. W. McKibben, Yue Hao, Gavin C. Conant, J. Chris Pires, and Michael S. Barker. 2021. “Genes Derived from Ancient Polyploidy Have Higher Genetic Diversity and Are Associated with Domestication in Brassica Rapa.” New Phytologist 230 (1): 372–86. 10.1111/nph.17194.

Reatini, Bryan, F. Alice Cang, Qiuyu Jiang, Michael T. W. McKibben, Michael S. Barker, Loren H. Rieseberg, and Katrina M. Dlugosch. 2022. “Data from: Chromosome-Scale Reference Genome and RAD-Based Genetic Map of Yellow Starthistle (Centaurea Solstitialis) Reveal Putative Structural Variation and QTLs Associated with Invader Traits.” Zenodo. 10.5281/zenodo.7324093.

Reatini, Bryan, and Todd J. Vision. 2020. “Genetic Architecture Influences When and How Hybridization Contributes to Colonization.” Evolution 74 (8): 1590–1602. 10.1111/evo.13972.

Reyes-Chin-Wo, Sebastian, Zhiwen Wang, Xinhua Yang, Alexander Kozik, Siwaret Arikit, Chi Song, Liangfeng Xia, et al. 2017. “Genome Assembly with in Vitro Proximity Ligation Data and Whole-Genome Triplication in Lettuce.” Nature Communications 8 (1): 14953. 10.1038/ncomms14953.

Rochette, Nicolas C., Angel G. Rivera-Colón, and Julian M. Catchen. 2019. “Stacks 2: Analytical Methods for Paired-End Sequencing Improve RADseq-Based Population Genomics.” Molecular Ecology 28 (21): 4737–54. 10.1111/mec.15253.

Ruan, Jue, and Heng Li. 2020. “Fast and Accurate Long-Read Assembly with Wtdbg2.” Nature Methods 17 (2): 155–58. 10.1038/s41592-019-0669-3.

Scaglione, Davide, Sebastian Reyes-Chin-Wo, Alberto Acquadro, Lutz Froenicke, Ezio Portis, Christopher Beitel, Matteo Tirone, et al. 2016. “The Genome Sequence of the Outbreeding Globe Artichoke Constructed de Novo Incorporating a Phase-Aware Low-Pass Sequencing Strategy of F1 Progeny.” Scientific Reports 6 (1): 19427. 10.1038/srep19427.

Stanke, Mario, Oliver Keller, Irfan Gunduz, Alec Hayes, Stephan Waack, and Burkhard Morgenstern. 2006. “AUGUSTUS: Ab Initio Prediction of Alternative Transcripts.” Nucleic Acids Research 34 (Web Server issue): W435–39. 10.1093/nar/gkl200.

Stepanova, Anna N., Joyce Robertson-Hoyt, Jeonga Yun, Larissa M. Benavente, De-Yu Xie, Karel Doležal, Alexandra Schlereth, Gerd Jürgens, and Jose M. Alonso. 2008. “TAA1-Mediated Auxin Biosynthesis Is Essential for Hormone Crosstalk and Plant Development.” Cell 133 (1): 177–91. 10.1016/j.cell.2008.01.047.

Tong, Chunfa, Huogen Li, Ying Wang, Xuran Li, Jiajia Ou, Deyuan Wang, Houxi Xu, et al. 2016. “Construction of High-Density Linkage Maps of Populus Deltoides × P. Simonii Using Restriction-Site Associated DNA Sequencing.” PLOS ONE 11 (3): e0150692. 10.1371/journal.pone.0150692.

Wang, Yupeng, Haibao Tang, Jeremy D. DeBarry, Xu Tan, Jingping Li, Xiyin Wang, Tae-ho Lee, et al. 2012. “MCScanX: A Toolkit for Detection and Evolutionary Analysis of Gene Synteny and Collinearity.” Nucleic Acids Research 40 (7): e49. 10.1093/nar/gkr1293.

Webb, David M., and Steven J. Knapp. 1990. “DNA Extraction from a Previously Recalcitrant Plant Genus.” Plant Molecular Biology Reporter 8 (3): 180–85. 10.1007/BF02669514.

Wendel, Jonathan F., Scott A. Jackson, Blake C. Meyers, and Rod A. Wing. 2016. “Evolution of Plant Genome Architecture.” Genome Biology 17 (1): 37. 10.1186/s13059-016-0908-1.

Widmer, Timothy L., Fatiha Guermache, Margarita Yu Dolgovskaia, and Sergey Ya. Reznik. 2007. “Enhanced Growth and Seed Properties in Introduced vs. Native Populations of Yellow Starthistle (Centaurea Solstitialis).” Weed Science 55 (5): 465–73.

Yang, Jian, Noah A. Zaitlen, Michael E. Goddard, Peter M. Visscher, and Alkes L. Price. 2014. “Advantages and Pitfalls in the Application of Mixed-Model Association Methods.” Nature Genetics 46 (2): 100–106. 10.1038/ng.2876.

