## Supplementary material for "Chromosome-scale reference genome and RAD-based genetic map of yellow starthistle (*Centaurea solstitialis*) reveal putative structural variation and QTL associated with invader traits": SI

**Appendix S1**

**Supplementary Information: “Chromosome-scale reference genome and RAD-based genetic map of yellow starthistle (*Centaurea solstitialis*) reveal putative structural variation and QTLs associated with invader traits”**

Bryan Reatini, Jessie A. Pelosi, F. Alice Cang, Qiuyu Jiang, Michael T. W. McKibben, Michael S. Barker, Loren H. Rieseberg, Katrina M. Dlugosch

**Supplementary Figures and Tables**

**Tables**

**Supplemental Table 1.** List of all genes identified as paleologs and their classification according to Frackify.

***STAR_Frackify.Class.csv***

**Supplemental Table 2.** List of candidate genes under QTL peaks and their identity as paleologs or non-paleologs.

***candidate_genes.csv***

**Supplemental Table 3.** Total number of nucleotide differences (top right triangle) and per-basepair rate of divergence (bottom left triangle) within gene regions for each pair of samples used in the gene annotation.

|  | **CAN066** | **CAN582** | **CAN665** | **GIL430** | **GIL559** | **GIL676** |
| --- | --- | --- | --- | --- | --- | --- |
| **CAN066** |  | 278257 | 278096 | 383501 | 345043 | 200500 |
| **CAN582** | 0.0046 |  | 305355 | 411176 | 366286 | 306978 |
| **CAN665** | 0.0046 | 0.0051 |  | 412930 | 365302 | 298919 |
| **GIL430** | 0.0064 | 0.0069 | 0.0069 |  | 385484 | 394206 |
| **GIL559** | 0.0058 | 0.0061 | 0.0061 | 0.006441 |  | 312353 |
| **GIL676** | 0.0034 | 0.0051 | 0.005 | 0.006587 | 0.005219 |  |

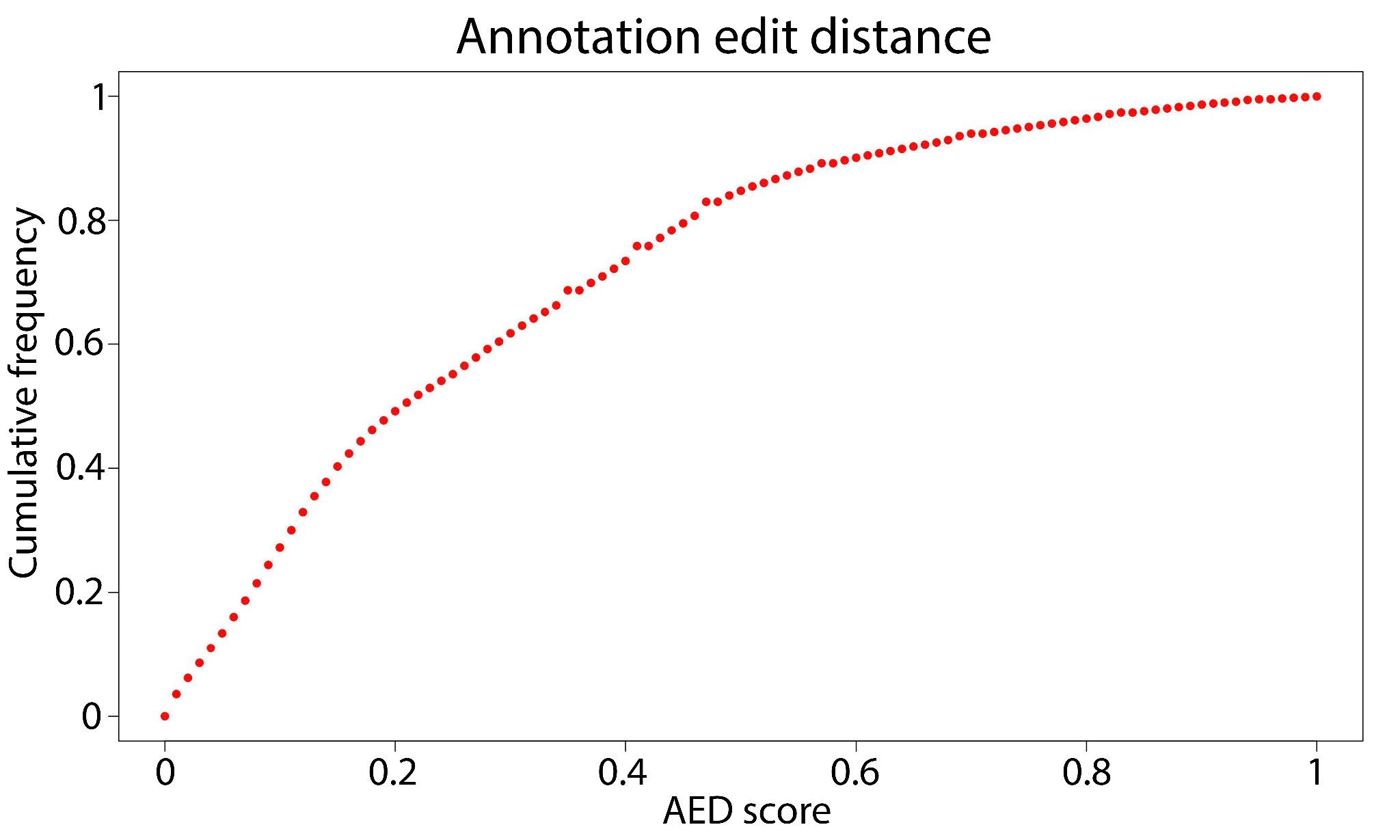

**Supplemental Figure 1**. Annotation Edit Distance (AED) for the *C. solstitialis* genome annotation.

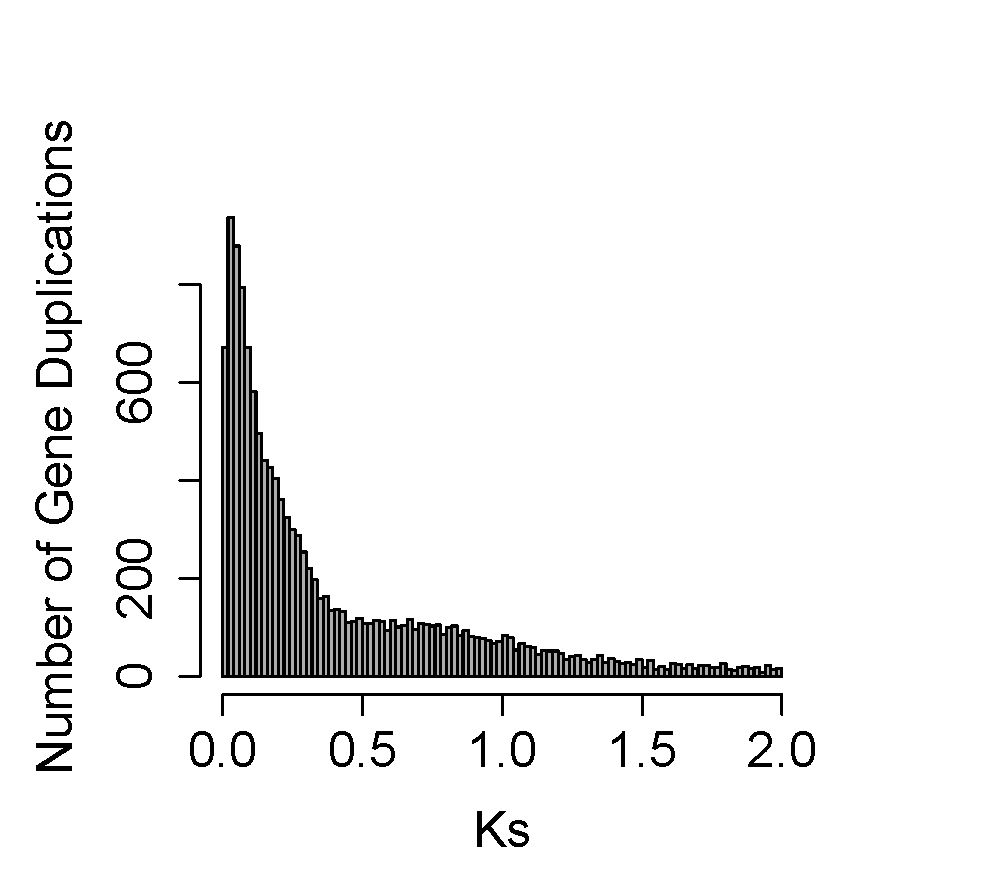

**Supplemental Figure 2**. Estimated gene duplications using DuPipe reveal a peak corresponding with the Compositae paleohexaploidy at Ks of 0.68.

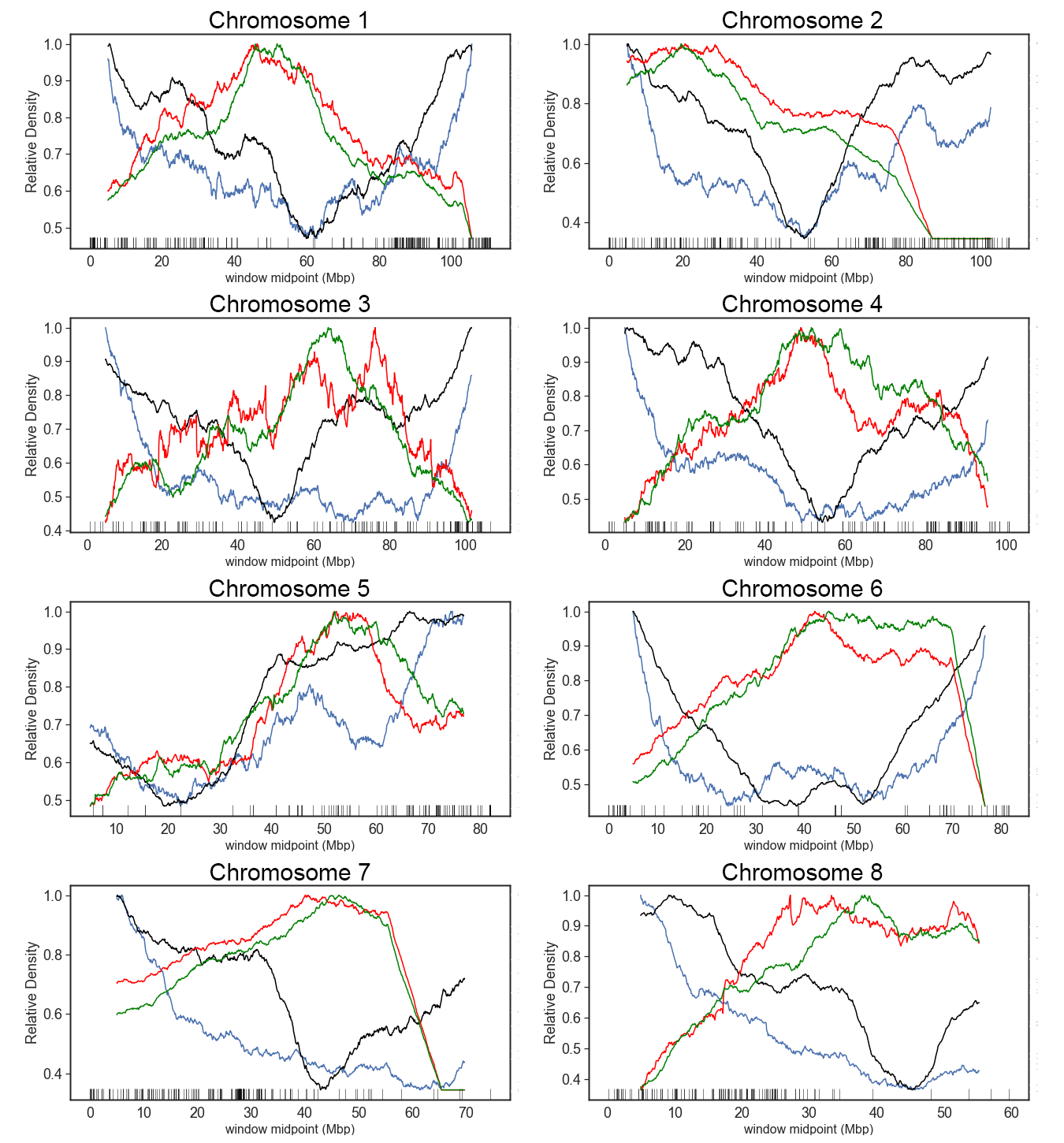

**Supplemental Figure 3**. Normalized density of genes (blue), unique repetitive elements (black), Gypsy LTR-RT elements (red), and Copia LTR-RT elements (green) across the genome. Rug plots show RAD marker density from the genetic map.

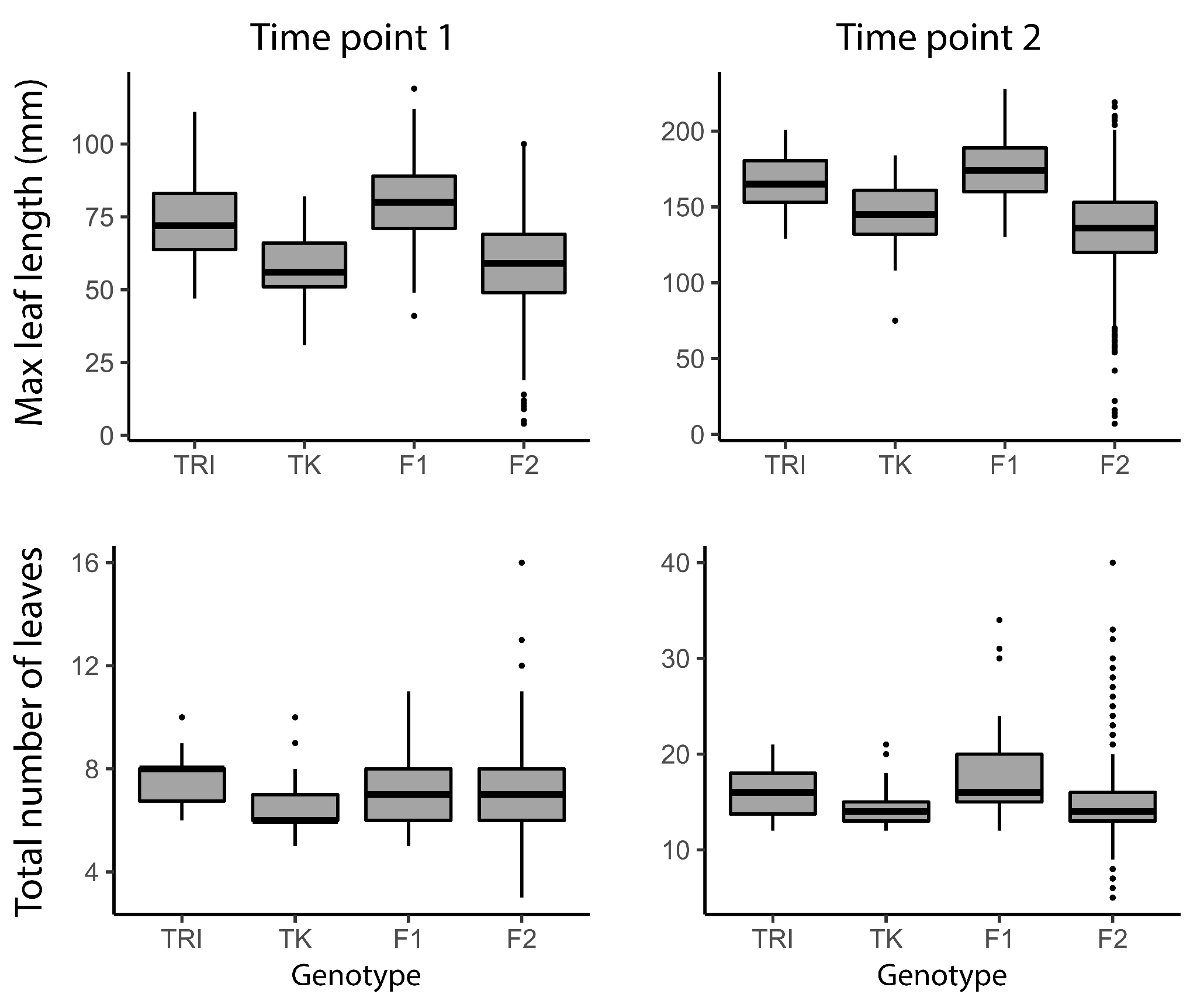

**Supplemental Figure 4**. Phenotypic values for invader parental genotype (TRI), native parental genotype (TK), F1, and F2 genotypes of the mapping population.

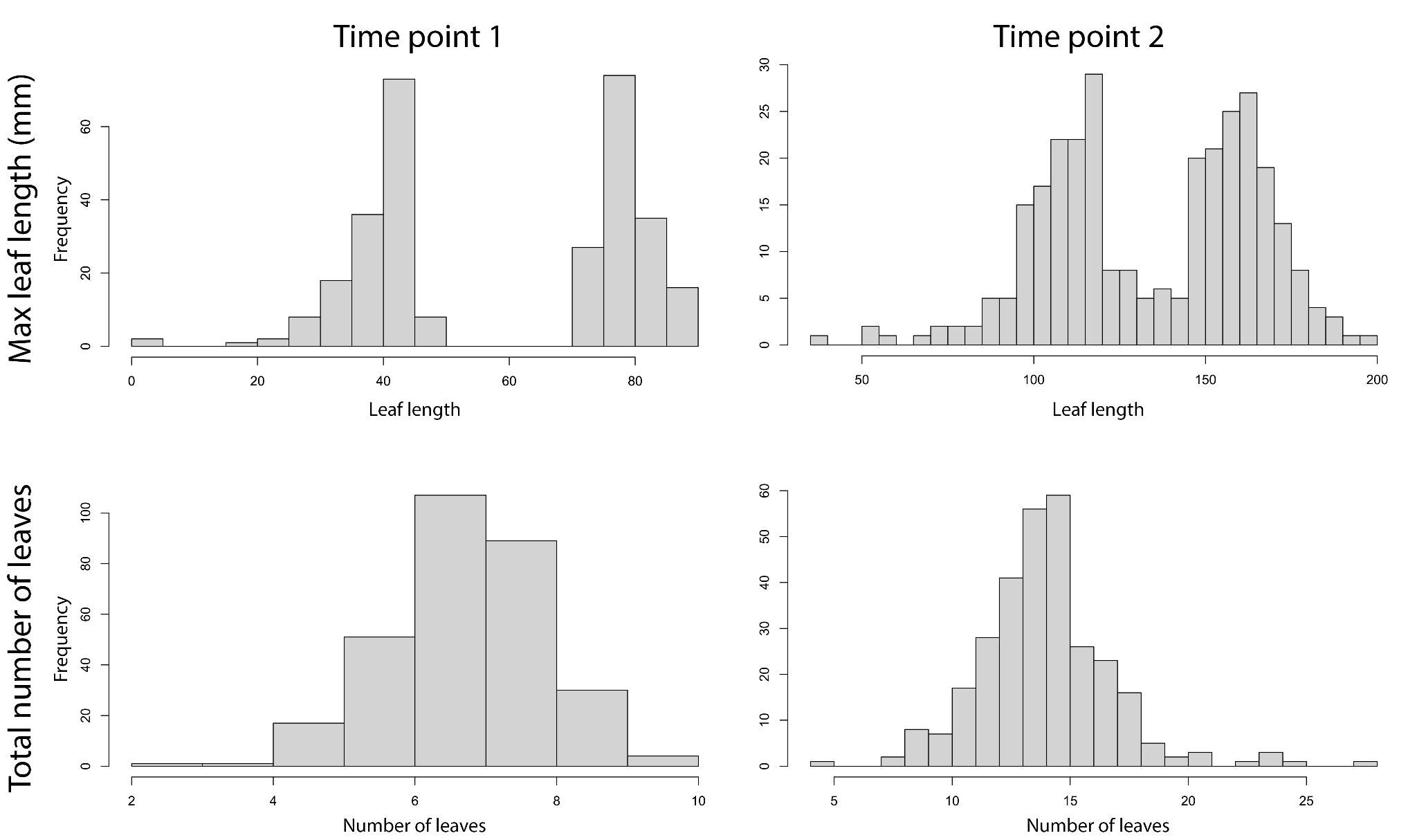

**Supplemental Figure 5**. Distribution of corrected values for maximum leaf length and total number of leaves at both time points for the 300 genotyped F2s used as input for QTL analysis.

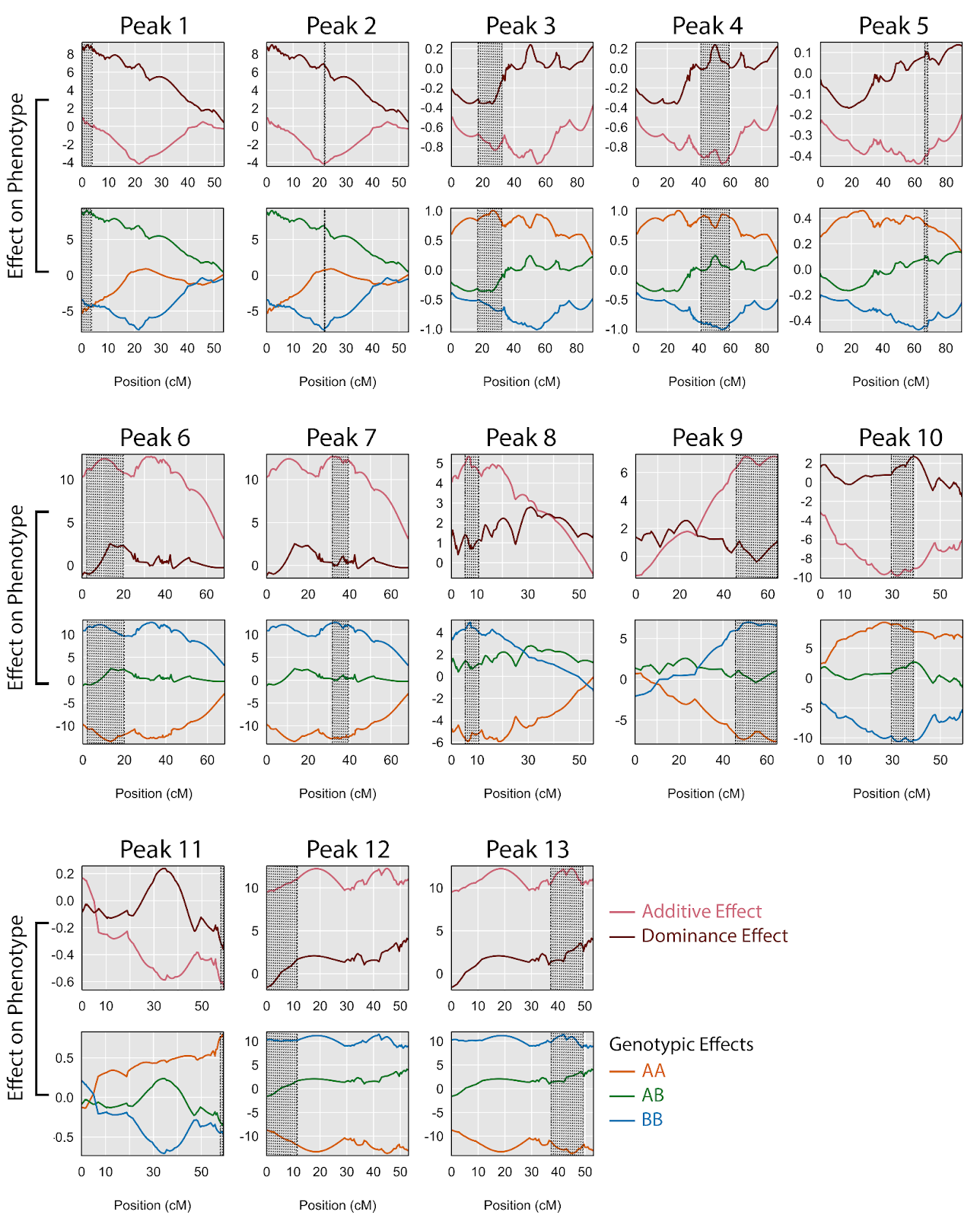

**Supplemental Figure 6**. Effect on phenotype for additive and dominance components (top) and genotypic effects (bottom) for each QTL peak summarized in Table 2. Effects are shown across the whole chromosome (in centimorgans) and the region of the QTL itself is shaded for each peak. For Genotypic effects, the A allele is the invader allele and the B allele is the native allele.  See Table 2 for a full description of the phenotypes and percentage of variation explained for each QTL peak.

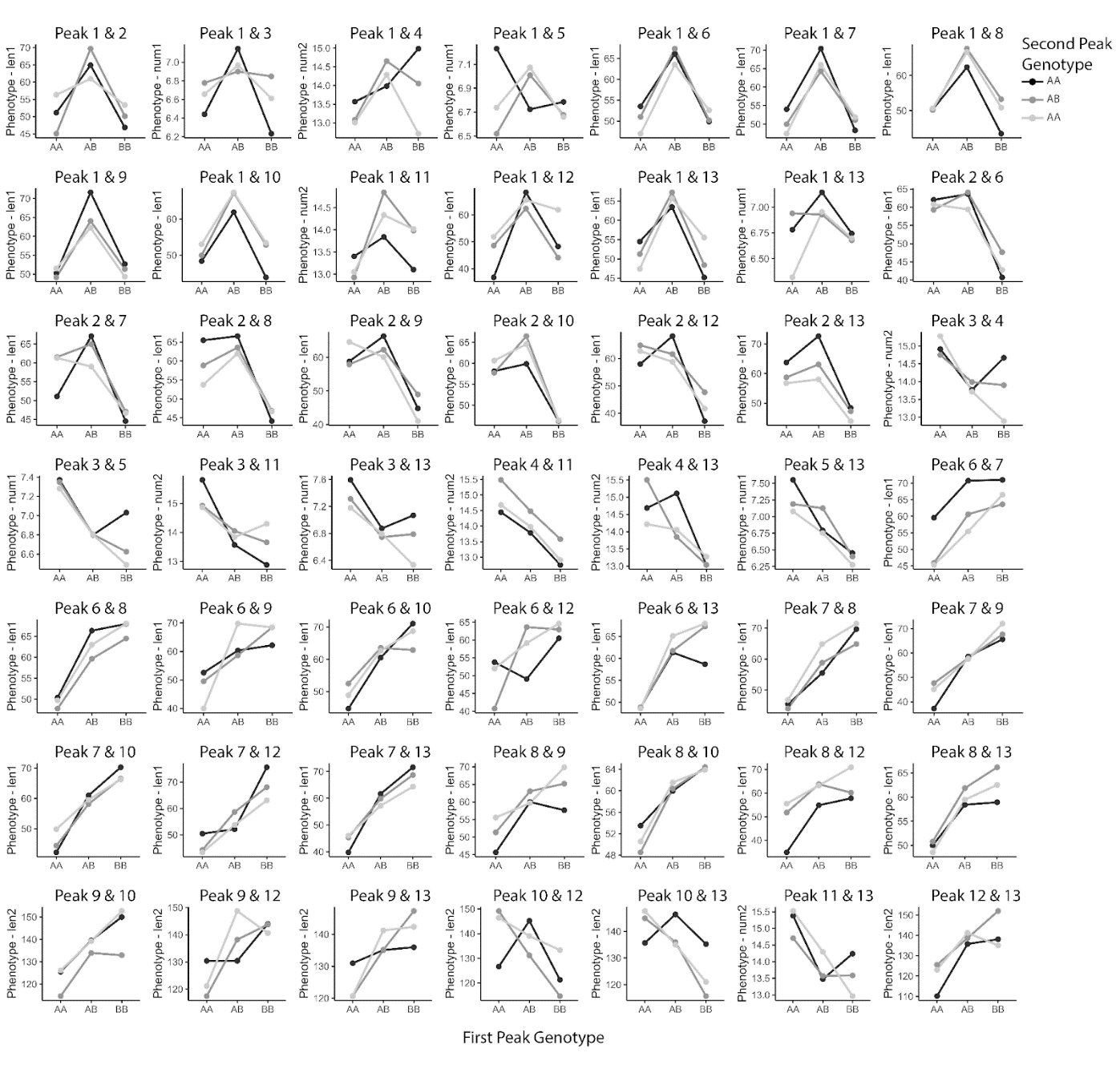

**Supplemental Figure 7**. Exploration of epistasis via pairwise interaction plots between the QTL associated with each phenotype. For each subplot, the genotype shown on the x-axis (the first peak genotype) is the genotype for the first QTL peak listed in the title and the genotype for the second QTL peak (shown by shaded lines) is the second peak listed in the title.

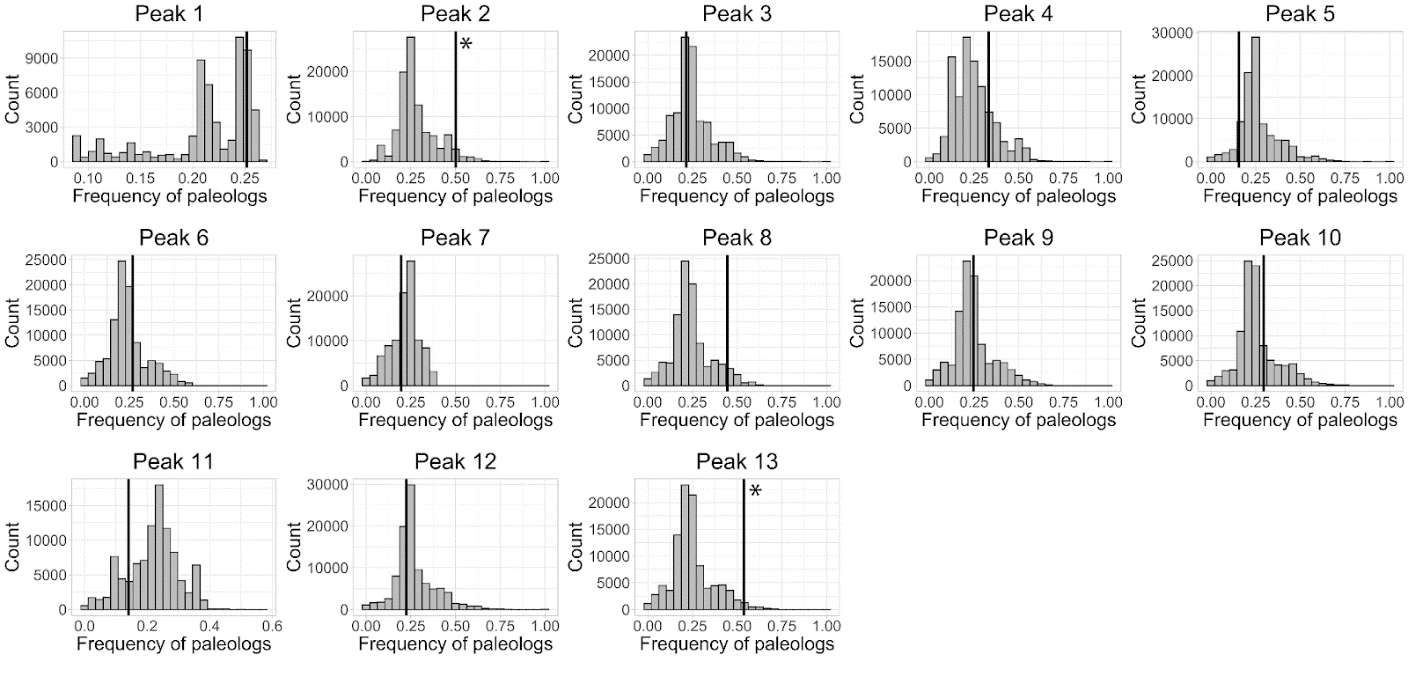

**Supplemental Figure 8**. Paleolog enrichment analysis using the null distribution of paleologs across the genome. Null distributions for 100,000 randomly selected blocks of the same size as each QTL peak are shown in gray histograms, and the corresponding frequency of paleologs within each QTL peak is marked by vertical black lines. Significant deviations from the null distribution are marked with asterisks.
